# Unleashing a Novel Function of Endonuclease G in Mitochondrial Genome Instability

**DOI:** 10.1101/2021.05.27.445952

**Authors:** Sumedha Dahal, Humaira Siddiqua, Shivangi Sharma, Ravi K. Babu, Meghana Manjunath, Sheetal Sharma, Bibha Choudhary, Sathees C. Raghavan

**Affiliations:** Department of Biochemistry, Indian Institute of Science, Bangalore 560012, India; Institute of Bioinformatics and Applied Biotechnology, Electronics City, Bangalore 560100, India; Department of Experimental Medicine, Postgraduate Institute of Medical Education and Research, Chandigarh, 160012, India

**Author notes:** To whom correspondence should be addressed: Department of Biochemistry, Indian Institute of Science, Bangalore-560012.

**Keywords:** Tetraplexes, Mitochondrial fragility, Endo G, Double-strand breaks, G4 DNA, MMEJ, Mitochondrial deletion

## Abstract

Having its own genome makes mitochondria a unique and semiautonomous organelle within cells. Mammalian mitochondrial DNA (mtDNA) is double-stranded closed circular molecule of about 16 kb coding 37 genes. Mutations, including deletions in the mitochondrial genome can culminate in different human diseases. Mapping of the deletion junctions suggests that the breakpoints are generally seen at hotspots. ‘9-bp deletion’ (8271-8281), seen in the intergenic region of cytochrome c oxidase II/tRNA^Lys^, is the most common mitochondrial deletion. While it is associated with several diseases like myopathy, dystonia, and hepatocellular carcinoma, it has also been used as an evolutionary marker. However, the mechanism responsible for its fragility is unclear. In the current study, we show that Endonuclease G, a mitochondrial nuclease responsible for nonspecific cleavage of nuclear DNA during apoptosis, can induce breaks at sequences associated with ‘9-bp deletion’, when it is present on a plasmid or in the mitochondrial genome. Through a series of *in vitro* and intracellular studies, we show that Endonuclease G binds to G-quadruplex structures formed at the hotspot and induces DNA breaks. Besides, we reconstitute the whole process of ‘9-bp deletion’ using purified Endonuclease G to induce breaks in mtDNA, followed by mitochondrial extract-mediated DSB repair and establish that microhomology-mediated end joining is responsible for the generation of mtDNA deletion. Finally, we show that the whole process is regulated by different stress conditions, which may modulate release of Endonuclease G to the mitochondrial matrix. Therefore, we uncover a new role for Endonuclease G in generating deletions, which is dependent on the formation of G4 DNA within the mitochondrial genome. Thus, in this study we identify a novel property of Endonuclease G, besides its role in apoptosis, and the recently described ‘elimination of paternal mitochondria during fertilization.

## INTRODUCTION

The mitochondria are semiautonomous organelle with their own genome. Mammalian mitochondrial genome encodes for 13 respiratory chain proteins, 22 tRNAs and 2 rRNAs (Chen and Butow, 2005; Clayton, 1984; Holthofer et al., 1999). Due to its vital role in oxidative phosphorylation, mitochondrial DNA is frequently exposed to reactive oxygen species and is more prone to DNA damage than nuclear DNA, leading to accumulation of 50 times more mutations, including deletions (Chen et al., 2011; Hudson et al., 1998; Michikawa et al., 1999; Pakendorf and Stoneking, 2005; Yakes and Van Houten, 1997). Mitochondrial mutations have been associated with aging and several disease conditions like myopathies, dystonia, cancer etc. (Chen et al., 2011; Taylor and Turnbull, 2005). These mutations include point mutations, mismatches, or deletions, which affect the coding of important proteins involved in oxidative phosphorylation and respiratory chain (Penta et al., 2001). Mitochondrial mutations, especially large-scale deletions of 200-5000 bp were shown to be associated with progression of breast, colorectal, renal, and gastric cancers (Bianchi et al., 2001; Carew and Huang, 2002; Modica- Napolitano et al., 2007; Penta et al., 2001). However, the correlation between mtDNA deletions and cancer remains unsettled. Three well studied phenotypes of mtDNA deletions are Kearns- Sayre Syndrome (KSS), Pearson syndrome, and progressive external ophthalmoplegia (PEO) (Marni, 2003). Large stretches of deletions seen in different diseased patients suggest the presence of fragile sites in the mitochondrial genome similar to the FRA sites of the nuclear DNA (Richards and Macaulay, 2001).

The FRA regions and other frequent chromosomal translocations have been observed in late replicating sites in the nuclear DNA, which are prone to replication fork stalling due to possible, stable secondary structure formation or inadvertent action of structure-specific nucleases (Dillon et al., 2010; Lin et al., 2013; Raghavan et al., 2004; Sun et al., 2001). Mitochondrial DNA has its own replication machinery that is not governed by cell cycle check points, and therefore, it is likely that the formation of a secondary structure may account for observed mitochondrial fragility.

Presence of alternate DNA structures like G-quadruplex, triplex, cruciform structures in mitochondrial genome have been reported along with their association with different mitochondrial diseases (Damas et al., 2012; Dong et al., 2014; Bharti et al., 2014; Oliveira et al., 2013). Breakpoint regions reported in case of PEO patients were at the proximity of G- quadruplex motifs, further suggesting the role of non-B DNA structures in mitochondrial disorders (Bharti et al., 2014). G4 forming motifs were also located adjacent to mitochondrial deletion breakpoints associated with KSS, a clinical subgroup of mitochondrial encephalomyopathies (Van Goethem et al., 2003; Zeviani et al., 1988). Further, a G4 ligand RHPS4 preferentially localizes to mitochondria and affects mitochondrial genome content, transcription elongation, and respiratory function (Falabella et al., 2019; Oliveira et al., 2013). Moreover, *in vitro* results suggested that a single-base mutation in mtDNA (mt10251) was sufficient to form G4 DNA (Chu et al., 2019). Importantly, ATP-dependent G4 resolving helicase Pif1 could unfold the non-B DNA and continue the synthesis by mitochondrial DNA polymerases, Pol γ and PrimPol (Butler et al., 2020). However, further studies are required to confirm the presence of these structures *in vivo* and establish their contribution towards mechanism of breakage and deletions.

Mitochondrial DNA (mtDNA) 9-bp deletion caused by loss of one copy of the 9-bp repeat sequence (CCCCCTCTA) in the intergenic region of cytochrome c oxidase II (COII)/tRNA^Lys^ was one of the most common deletions in mitochondria (Redd et al., 1995; Thomas et al., 1998). It has been associated with various diseases including hepatocellular carcinoma (Jin et al., 2012; Komandur et al., 2011; Krishnan and Turnbull, 2010; Zhuo et al., 2010). The 9-bp deletion was also widely used as a phylogenetic marker to study evolutionary trends and migration of populations (Soodyall et al., 1996; Yao et al., 2000). The occurrence of deletions in such high frequency indicates the fragility of the region; however, its mechanism is unclear.

Out of the large scale mtDNA deletions reported, one of the well-studied examples is mt^4977^, and accumulation of it results in different mitochondrial disorders including cancer (Yusoff et al., 2019). mt^4977^ eliminates mitochondrial sequences between 8470 and 13447 bp, which include removal of genes required for normal oxidative phosphorylation (OXPHOS) (Lee et al., 1994; Yang et al., 1994; Yen et al., 1991). Recently the role of replication slippage and DSB repair has been shown to play a role in the generation of mtDNA^4977^ in mammalian cells (Phillips et al., 2017). Previous studies have shown that presence of a 13 bp direct repeat play a major role in mt^4977^, as only one of the repeats was retained following joining indicating the involvement of microhomology mediated end joining (Samuels et al., 2004; Srivastava and Moraes, 2005). When restriction endonucleases were used to introduce DSBs, there were accumulation of deletions, which further implicates the imperfect repair and deletion in the mitochondrial genome (Srivastava and Moraes, 2005). A previous study from our laboratory suggested the existence of MMEJ in mitochondria, which operate in a microhomology dependent manner explaining deletions frequently seen in mitochondrial DNA (Tadi et al., 2016).

Here we investigate the mechanism of fragility associated with the most common deletion seen in the mitochondrial genome. The putative role of non-B DNA structures, mechanism of DNA breakage and repair associated with the formation of mtDNA deletions are extensively studied. We report that the region associated with 9-bp deletion can fold into G-quadruplex DNA structure using various biochemical and *ex vivo* methods. For the first time we show that Endonuclease G can specifically bind to G-quadruplex structure and cleave the mtDNA. Finally, the microhomology mediated repair of the broken DNA leads to the formation of “9-bp deletion” seen in human mitochondria. The whole process is controlled by different stress conditions, which may regulate the release of Endonuclease G to the mitochondrial matrix.

## RESULTS

Previous studies have reported different non-B DNA motifs in mitochondrial genome and suggested such alternate form of DNA may play a role in deletions in mitochondria (Damas et al., 2012; Dong et al., 2014). *In silico* studies revealed that 5 classical G4 DNA motifs present in mitochondrial genome could fold into G4 DNA (Cer et al., 2013). Since “9-bp deletion” (8271- 8281) is the most frequent mitochondrial deletion and occurs next to one of the 5 G4 motifs with the coordinate position 8252-8295 (designated as mitochondrial Region I), we have investigated putative role of G4 DNA structure and mechanism of DNA breakage and repair associated with the formation of the mtDNA deletions. Importantly, the chosen G-quadruplex motif was also present in other 2 large scale deletions.

### G-quadruplex motifs from Region I of mitochondrial DNA, when present on shorter DNA or on a plasmid DNA can fold into G-quadruplex structure

To investigate whether mitochondrial Region I can indeed form G-quadruplex structures, oligomeric DNA containing the G-rich region referred to as “G1” and complementary C strand (C1), were synthesized (Figure 1A). Radiolabelled oligomeric DNA were then subjected to electrophoretic mobility shift assay in the absence or presence of KCl. Faster mobility of G-rich strand compared to its complementary C-rich strand suggested the formation of both intra and intermolecular G4 DNA structure in a KCl dependent manner (Figure 1B). Importantly, formation of intramolecular G4 DNA was abrogated when G-stretches were mutated (Figure 1A; S1A). Further, circular dichroism (CD) studies revealed that the spectra exhibited characteristic of a parallel G-quadruplex DNA with absorption maxima of 260-270 nm and absorption minima of 240 nm as opposed to the C-rich strand, which showed the maxima similar to that of a typical single-stranded DNA (Figure 1C).

**Figure 1.**
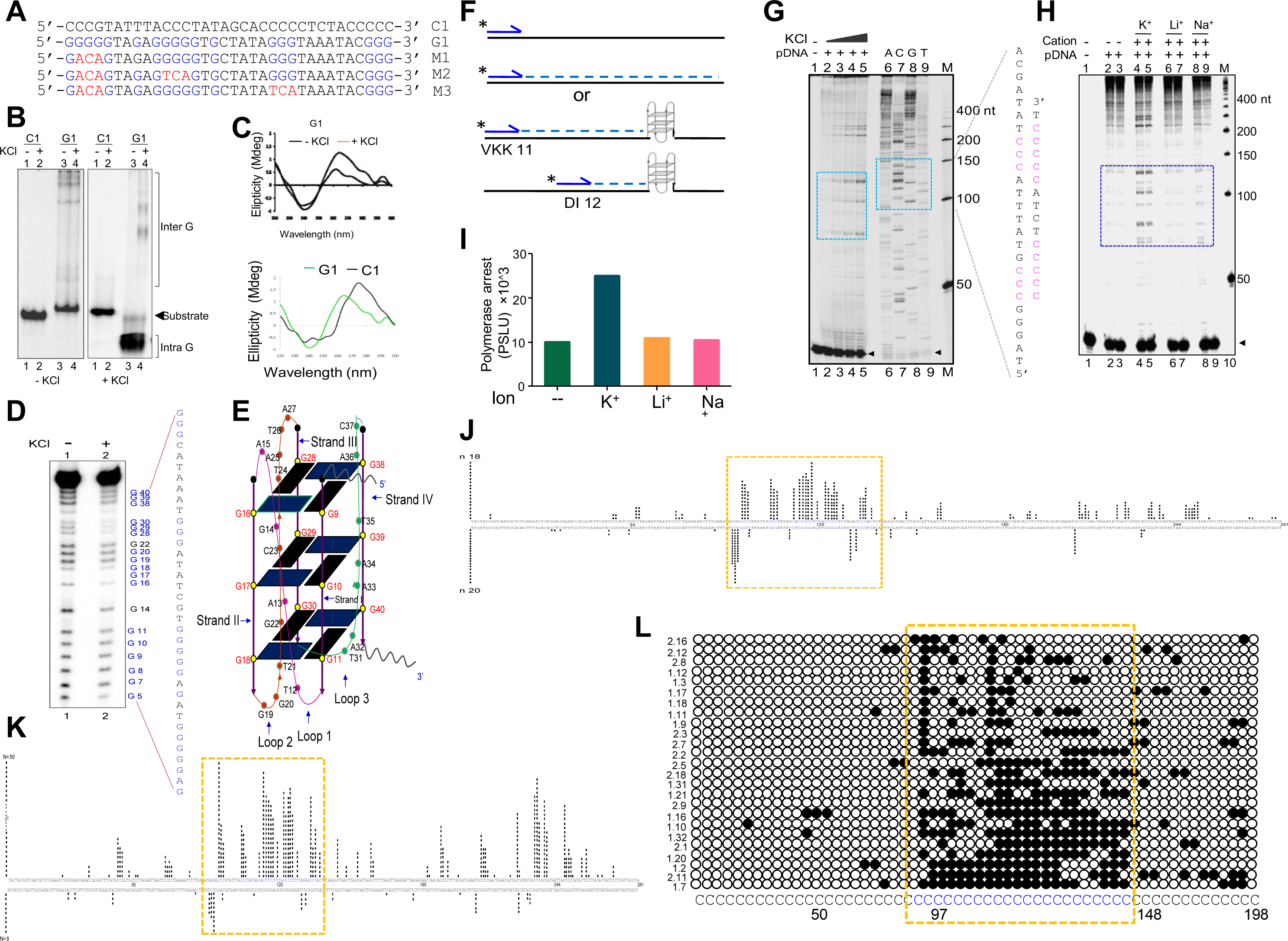
Biochemical studies to investigate formation of G-quadruplex structure in Region I of mitochondrial genome. **A.** The oligomeric sequence of the G-rich strand (indicated by ‘G1’), and the complementary C-rich strand (indicated by ‘C1’) are derived from the predicted mitochondrial G-quadruplex forming Region I. The stretches of guanines are marked in blue and mutations to G-stretch in red. **B.** The G and C-rich strands were incubated in presence of 100 mM KCl and resolved on 15% native polyacrylamide gels in the absence or presence of KCl (100 mM), in the gel and running buffer. The substrate, intramolecular (Intra G), and intermolecular (Inter G) quadruplex structures are indicated. **C.** CD spectra for G1 in presence (red) or absence of 100 mM KCl (black). CD spectra for G1 (green) and C1 (black) in the presence of 100 mM KCl. In each case where KCl was added, the respective oligomer resuspended in Tris-EDTA buffer was incubated for 1 h at 37°C and the spectra was recorded using JASCO J-810 spectropolarimeter (scan range of 220-300 nm). **D.** DMS protection assay for the Region I of mitochondria. The oligomer DNA was allowed to fold into G4 DNA and then treated with DMS, followed by cleavage with piperidine. The products were resolved on a 15% denaturing PAGE. Substrate indicates the gel-purified DNA from the reaction incubated with or without KCl. All the positions of guanines are indicated. **E.** A representative 2D model for intramolecular G-quadruplex structure formed at mitochondrial Region I based on the reactivity of guanine to DMS. **F.** Schematic showing the primer extension across plasmid DNA containing a mitochondrial Region I. Positions of primers, VKK11 and DI12 are indicated. The radiolabeled primer binds to one of the strands of the plasmid DNA upon heat denaturation and extends till it encounters a non-B DNA, as it blocks the progression of the polymerase. The products (dotted lines) are then resolved on a denaturing PAGE. **G.** The plasmid, pDI1, containing the mitochondrial Region I was used for primer extension studies using radiolabeled primers for the G-rich (VKK11) in a KCl dependent manner. Sequencing ladder was prepared, using VKK11 by the chain termination method of sequencing. Pause sites are shown with dotted rectangles. Sequence corresponding to the pause site is boxed and G-quadruplex forming motifs are indicated. “M” denotes the 50 nt ladder. **H.** Evaluation of the effect of different cations (Na^+^, K^+^, Li^+^) on non-B DNA formation at mitochondrial Region I by using primer extension assay. The pause sites are indicated in a rectangular box. **I.** Bar diagram showing the effect of ions in primer extension assay. The values are expressed in PSL units representing the extent of pause. **J.** Bisulphite modification assay on plasmid (pDI1) containing mitochondrial Region I. Vertical bars represent the number of times the respective cytosine in the top and bottom strands of mitochondrial Region I is converted to thymine after deamination when treated with sodium bisulphite, followed by PCR. **K, L.** Bisulphite sequencing on the mitochondrial genome for determining the formation of G4 DNA at Region I. Each vertical bar represents the conversion of a cytosine to uracil in the top strand or bottom strand. A total of 59 clones were sequenced from both top and bottom strands (K). Single strandedness observed in a DNA fragment of 198 nt containing Region I of mitochondrial genome following bisulphite modification assay (L). Each row represents cytosines present in a DNA molecule. Each dark circle represents the conversion of a cytosine to thymine on top strand after deamination upon treatment of sodium bisulphite, followed by PCR and DNA sequencing. Of the 59 clones sequenced, most reactive 25 molecules are shown (L). Sequences at the G-quadruplex forming motif are shown in blue and boxed among the molecules. Refer also Figure S1E.

DMS protection assay was performed to understand the precise base pairing of guanine nucleotides involved in the G4 plates compared to the ones, which lie within connecting loops that facilitate DNA strand folding and thus G-quadruplex formation (Nambiar et al., 2011). DMS methylates the N7 position of the guanine base, making it susceptible to subsequent piperidine cleavage. Since the N7 position of G residues forming the G4 plate is engaged upon Hoogsteen base pairing, these residues are protected from DMS treatment and piperidine cleavage. When the G-quadruplex sequences from mitochondrial Region I (G1) was subjected to DMS protection assay, following structure formation, we observed that the majority of G residues in this region showed protection, suggesting that they were indeed involved in Hoogsteen bonding (Figure 1D, lane 2). The results, therefore, demonstrate that the guanine residues from the G stretches of mitochondrial Region I motif participate in G-quadruplex formation. Based on this and other information from above studies, a parallel, intramolecular G-quadruplex structure was modelled (Figure 1E). Thus, our studies including the DMS modification assay reinforces that the Region I of mitochondrial DNA can fold into intramolecular G-quadruplex (Figure 1E).

In the context of double-stranded DNA, G-quadruplexes can form only when the Hoogsteen base pairing between the guanine residues is stronger than the Watson and Crick H- bonding required for duplex B-DNA formation. Therefore, the G-quadruplex forming ability of the mitochondrial Region I was tested after cloning into a plasmid, pDI1. Interestingly, an inverted repeat (coordinate positions 8295-8363), which could fold into a cruciform DNA was also present adjacent to the G4 motif, thus that was also part of the pDI1. Further, to delineate the role of G4 DNA in the plasmid context, a mutant plasmid with scrambled sequence corresponding to G4 motif but with intact inverted repeats (pDI2) was also constructed (Figure S1B). Previous studies have shown that the formation of G4 DNA can lead to a pause during replication (Kumari et al., 2015; Nambiar et al., 2013; Voineagu et al., 2008; Wells, 2009). To assess whether the G4 DNA formed at Region I of mitochondrial genome can act as a replication pause, primer extension assay was performed using pDI1 (Kumari et al., 2015) (Figure 1F). A sequencing ladder was also generated for mapping the location of the polymerase pause sites on the plasmid (Figure 1G). Results revealed multiple polymerase pause sites and the intensity of the pauses increased with addition of higher concentrations of KCl (Figure 1G). Interestingly, there was no pause observed in the absence of KCl, indicating the absence of formation of intramolecular G-quadruplex species in these cases (Figure 1G, lane 2). Considering that formation of cruciform DNA occurs irrespective of the presence of KCl, result also suggest that the observed pause was due to formation of G4 DNA, but not cruciform DNA. Besides, the observed polymerase pauses mapped to the G-quadruplex motif (Figure 1G, lanes 7-10). Further, primer extension assays using a nested primer approach (Figure 1F) showed that the primer VKK11 that annealed farther away from the G-quadruplex motif gave longer polymerase stop products (Figure S1C, lanes 2-3) compared to primer DI12, that was positioned closer (Figure S1C, lanes 5-6) in a K^+^ dependent manner. It was also observed that the appearance of strong polymerase pause sites were dependent only on K^+^ (Figure 1H, lanes 4-5; I), but not on other monovalent cations such as Li^+^ (with no or little G-quadruplex stabilizing properties) or Na^+^ (with limited stabilizing properties) (Figure 1H, lanes 6-9; I) suggesting that the non-B DNA that is responsible for polymerase arrest was G4 DNA. Finally, abrogation of G4 DNA structure by mutation of G4 DNA motif (pID2) resulted in disappearance of the observed pause site further confirming above results (Figure S1D).

### Mitochondrial Region I can exist as G-quadruplex structure within the mitochondrial genome and when present as part of a plasmid DNA

Formation of G4 DNA on a plasmid or mitochondrial genome could result in single-strandedness at complementary and at adjacent regions corresponding to G4 motif. Sodium bisulphite modification assay was performed on pDI1 to check the single-strandedness in the complementary C strand of Region I. Sodium bisulphite deaminates cytosine resulting into uracil, when present on single-stranded DNA, and this change can be read as a C→T conversion after PCR and sequencing (Figure S1E). Out of a total of 40 clones sequenced, 38 molecules showed C to T conversion to varying extents in either top or bottom strands (Figure 1J). Interestingly, the cytosines complementary to the G-quadruplex motif exhibited single-stranded nature (Figure 1J), indicating the formation of a G-quadruplex structure in the G-rich strand. Importantly, ∼15% molecules exhibited continuous conversion at the region corresponding to the G4 motif. These results suggest that Region I of mitochondrial DNA can fold into a non-B DNA even when cloned into a plasmid.

Further, to investigate existence of G4 DNA structures in the mitochondrial genome, sodium bisulphite modification assay was performed after extracting the mitochondrial genome from human cells (Nalm6) under nondenaturing conditions. Results showed that out of 59 DNA molecules analyzed following cloning and sequencing, 52 showed C→T conversions to varying extents (Figures 1K, L). Cumulative analysis of converted cytosines on a fragment of 281 bp revealed single-strandedness at Region I corresponding to complementary region of G4 motifs. When the mitochondrial genome was analyzed, regions adjacent to the G4 motif also showed certain levels of single-strandedness (Figure 1K). However, C→T conversion rate was significantly lower when upstream and downstream sequences were analyzed for conversion. Further, among the 50 clones from the top strand, 14 of them (28%) had continuous conversions of 11 or more cytosines when a stretch of 20 cytosines were considered on a single DNA molecule level (Figure 1L). This suggests that the formation of G4 DNA at Region I may not occur in all DNA molecules of the mitochondrial genome at a given time. However, it is evident that the G-quadruplex motif at Region I can indeed fold into G-quadruplex structure in mitochondria.

### BG4, a G-quadruplex binding antibody binds to different G-quadruplex structures in the mitochondrial genome

From the previous studies, it was established that the BG4 antibody has the ability to bind to G-quadruplexes (Chambers et al., 2015; Das et al., 2016; Javadekar et al., 2020). In order to study whether BG4 indeed binds to mitochondrial genome inside cells, immunofluorescence in HeLa and 293T cells was performed after testing the specificity of antibody (Javadekar et al., 2020). Either Mito-tracker Deep Red or Mito-tracker Green FM was used for staining mitochondria while the nucleus was stained with DAPI. BG4 was stained with Alexa fluor tagged secondary antibodies. Merging of green and red foci resulting in yellow foci was observed in the case of both HeLa and 293T cells, which revealed localization of BG4 to mitochondria (Figures 2A, S2A). The colocalization was further analysed and confirmed using JaCoP plugin in ImageJ software. In both cases, an image with Coste’s mask is presented, which shows the extent of localization in the form of a white dot after the merge of red and green foci (Figures 2A-C). Colocalization analyses show Mander coefficient value between 0.25-0.4, irrespective of the stain used as MitoTracker, in case of HeLa and 293T cells suggesting that G-quadruplex structures can be detected in the mitochondrial genome in cellular context (Figure 2B, C). To further confirm that the colocalization was indeed between the MitoTracker and BG4 antibody, immunofluorescence was performed in rho(0) cells (human osteosarcoma cells with depleted mtDNA) (Dong et al., 2017; Tan et al., 2015). Results suggested that although BG4 was observed in the cytoplasm of rho(0) cells, they do not colocalize with mitochondria (Figure 2D, E).

**Figure 2.**
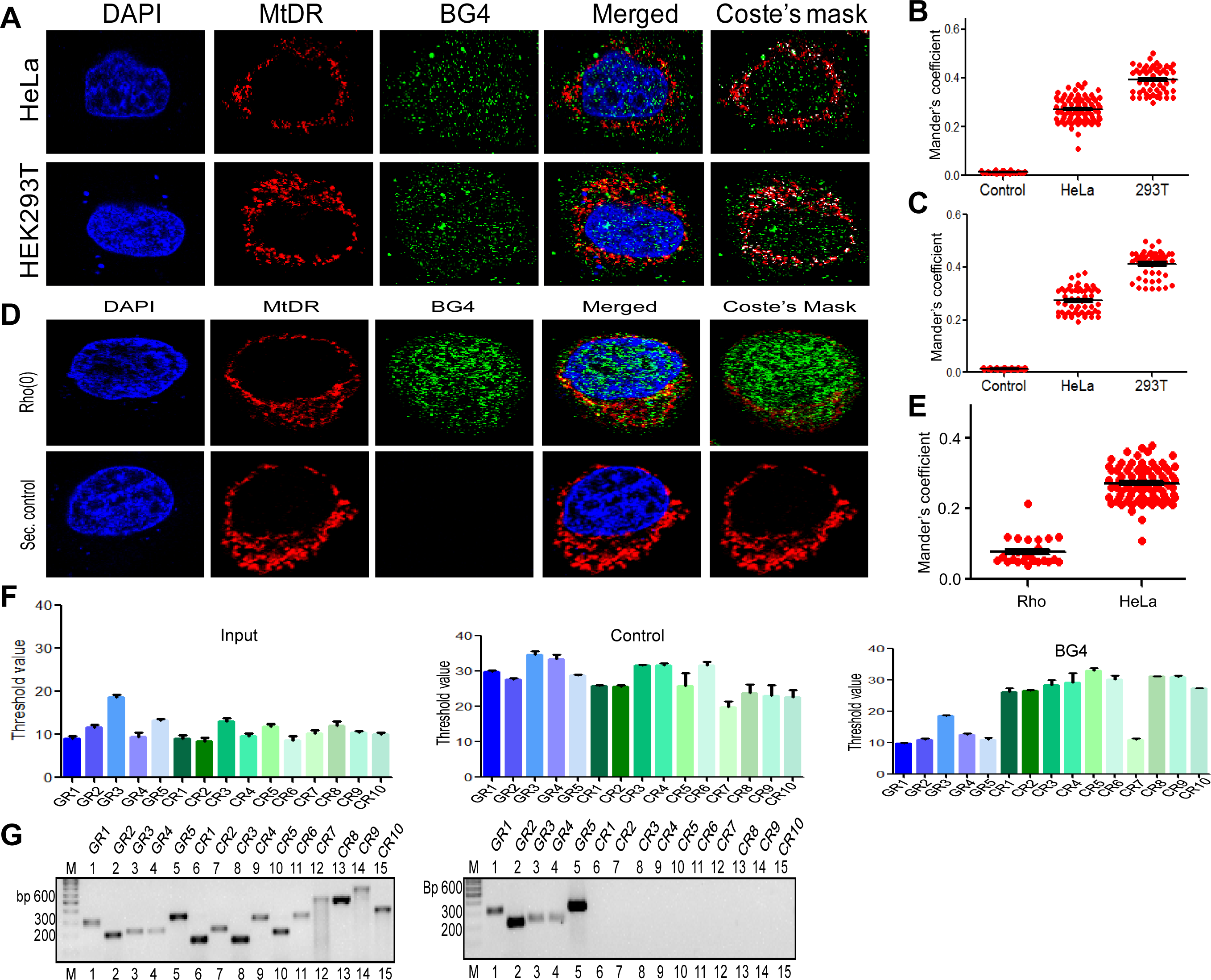
Evaluation of G-quadruplex structure in mitochondrial genome within cells. **A.** Representative images of HeLa and HEK293T cells showing localization of BG4, the G4 binding antibody to mitochondria following immunofluorescence study. Nucleus is stained with DAPI (blue color), mitochondria with MitoTracker DR (red) and BG4 with Alexa-Fluor 488 (green). A merged image is shown with merge of red and green as depicted by Coste’s mask (colocalization is represented as a white dot). **B, C.** Quantitation showing colocalization of BG4 with MitoTracker indicated as dot plots. The colocalization was quantified using Mander’s colocalization coefficient (ImageJ software) analyzing a minimum of 100 cells as red over green (B) and green over red (C). **D.** Representative image of rho(0) cell showing localization of BG4 to mitochondria following immunofluorescence as investigated in panel A. **E.** Quantitation showing comparison of colocalization of BG4 between HeLa cells and rho(0) cells is shown as dot plots. The colocalization was quantified using Mander’s colocalization coefficient analyzing a minimum of 50 cells as red over green. **F.** BG4 bound mtDNA was purified after reverse crosslinking and used for real-time PCR using primers derived from different regions of the mitochondrial genome, which include 5 G-quadruplex forming regions and 10 random regions. Input DNA served as template control. No antibody control was also used. Bars in blue (first 5) are for G-quadruplex forming regions, while in green (last 10) are for random regions. Y-axis depicts threshold Ct value obtained following real time PCR for each primer. Error bar represents mean ± SEM. **G.** Agarose gel profile showing the amplification of Input DNA (left panel) and BG4 pull down DNA (right panel). “M” denotes 100 bp ladder.

To examine whether BG4 can indeed bind to mitochondrial G-quadruplexes, *in vitro* binding of purified BG4 to oligomers harbouring G-quadruplex region was performed (Figure S2B). Results suggested that the bound complex was seen when G-rich oligomer (G1) was incubated, while such binding was absent when incubated with C-rich oligomer (C1) (Figure S2B).

To investigate whether BG4 can bind to G-quadruplex DNA present in the mitochondrial genome, ChIP-qPCR strategy was used (Figures S2C, D). Briefly, following crosslinking, mitochondria were isolated and region binding to BG4 was immunoprecipitated. Five regions corresponding to G4 motifs and ten random regions without G4 motifs were selected for analysis by using appropriate PCR primers (Figure S2D). Results showed that all five G-quadruplex forming regions showed amplification between 10-18 cycles when qPCR was performed (Figure 2F). In contrast, 9 of 10 random regions did not show any amplification following BG4 pull down (Figure 2F). Samples with input DNA served as a positive control and as expected, all the regions were amplified. Samples where no antibody was added, did not result in amplification, when G4 specific or random region primers were used revealing the specificity of BG4 towards G4 DNA (Figure 2F). Similar results were also obtained when BG4 pull down samples were evaluated using semiquantitative PCR (Figure 2G). However, none of the regions amplified when samples from secondary control was used (Figure 2F). Sequence analysis showed that the control region that was amplified following BG4 pull down had high GC content, although no G4 motifs were seen and require further analysis (Figure 2F).

Thus, our results suggest that Region I and other G4 motifs in the human mitochondrial genome can indeed fold into G-quadruplex DNA. However, based on bisulphite DNA sequencing from Region I, it appears that structure may not be formed in all DNA molecules, and the frequency may be <25%.

### Mitochondrial extract induces cleavage at or near G-quadruplex structure at Region I

In order to elucidate the mechanism responsible for cleavage at mitochondrial Region I, the involvement of mitochondrial nucleases was considered. To test this, mitochondrial extract was isolated from the rat tissues (testes and spleen) and used for cleavage assay. The purity of the extract was confirmed by western blotting using the anti-Cytochrome C (mitochondrial marker), anti-PCNA (nuclear or cytosolic marker), and anti-Actin (loading control) (Figure 3A). Plasmid harbouring mitochondrial Region I (pDI1) was treated with the mitochondrial extract (0.25, 0.5, 1 and 2 μg) and resolved on an agarose gel following deproteinization. Interestingly, a concentration dependent increase in cleavage efficiency (supercoiled form to nicked form) was observed, and the DNA nicking was only one site per plasmid molecule (Figure 3B, lanes 1-5). To test whether the cleavage observed was due to the presence of G4 DNA, the plasmid bearing mutation in mitochondrial Region I and intact adjacent inverted repeats (pDI2) was treated with mitochondrial extract and analysed. Interestingly, there was a significant reduction in cleavage when two stretches of G’s were mutated (Figure 3B, lanes 6-10) indicating the cleavage was indeed due to the presence of G quadruplex and not due to inverted repeats.

**Figure 3.**
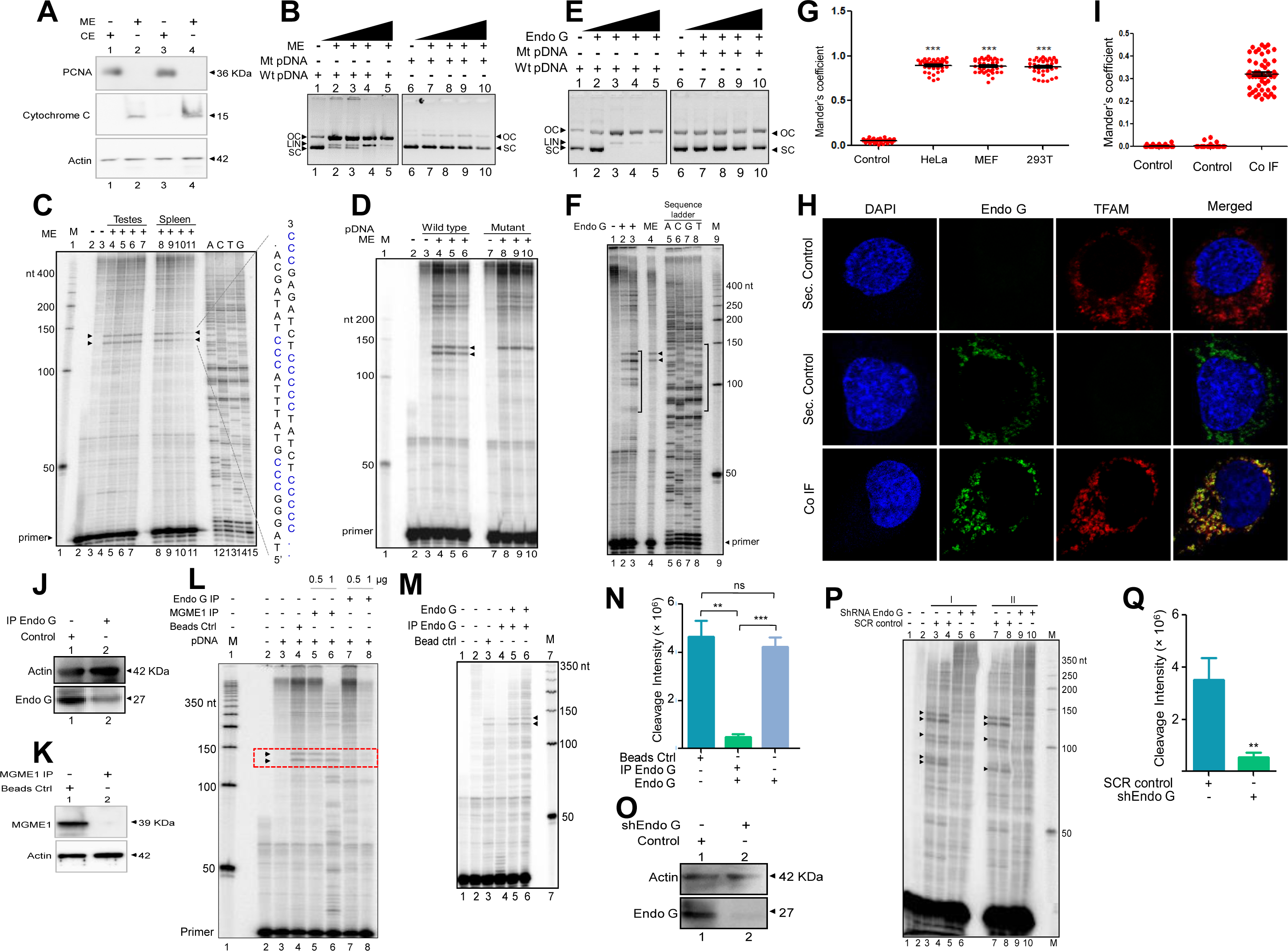
Studies to identify mitochondrial nuclease responsible for cleavage at G4 DNA formed at Region I of the mitochondrial genome. **A.** Purification and characterization of mitochondrial extracts. Mitochondrial extract was prepared from rat testis (RT) and spleen (RS), and its purity was evaluated using specific markers by western blotting. Purity of CE (cytosolic extract) and ME (mitochondrial extract) were determined using antibodies against PCNA (nuclear marker), cytochrome C (mitochondrial marker) and Actin (loading control). B. *In vitro* nicking assay on a plasmid containing wild type (WTpDNA) mitochondrial Region I (pDI1) and mutant (MtpDNA) of Region I (pDI2). Both wild type and mutant plasmids were incubated with mitochondrial extract from rat testes (RT-ME) and resolved on a 0.8% agarose gel. Lanes 1 and 6 served as the control without any extract in the reaction, whereas lanes 2-5 and lanes 7-10 are with increasing concentration (0.5, 1, 1.5 and 2 µg) of mitochondrial extract for pDI1 and pDI2, respectively. **C.** Primer extension assay on plasmid containing mitochondrial Region I (pDI1) to determine the position and location of cleavage. pDI1 was incubated with either ME prepared from testes or spleen, reaction products were purified and used for primer extension assay. Pause sites are indicated, which correspond to G4 DNA motif. Sequencing ladder was used as marker. **D.** Primer extension assay on pDI1 and pDI2 (G4 mutant) following incubation with RT-ME. E. *In vitro* nicking assay using purified Endonuclease G on wild type and mutant plasmids containing mitochondrial Region I (pDI1 and pDI2). Both wild type and mutant plasmids were treated with increasing concentration of purified Endonuclease G and resolved on a 0.8% agarose gel. Lanes 1 and 6 served as the control without any protein in the reaction whereas lanes 2-5 and lanes 7-10 are with increasing concentration (30, 60, 90, 120 ng) of purified protein. **F.** Cleavage assay was performed on a plasmid containing mitochondrial Region I, pDI1 following incubation with purified Endonuclease G. Primer extension assay was carried out using γ-^32^P radiolabeled VKK11 primer and resolved on 8% denaturing polyacrylamide gel. Lane 1 is without protein, lane 4 is with mitochondrial extract and lanes A, C, G, T represents the sequencing ladder. Lanes 2 and 3 represent 30 and 60 ng of Endonuclease G incubated samples. ‘M’ is 50 bp ladder. **G.** Colocalization analyses of Endonuclease G and Mitotracker signals using JaCoP in ImageJ software based on immunofluorescence studies performed in multiple cell lines (see Figure S4C). Minimum of 50 cells were used for analysis of colocalization of red and green signals and plotted as the colocalization value of green overlapping red. *Y-*axis depicts the Mander’s colocalization coefficient value calculated for green over red and plotted in the form of dot plot. The significance was calculated using GraphPad Prism 5.0 with respect to secondary control and shown as mean ± SEM (*ns* not significant, \**p* < 0.05, \*\**p* < 0.005, \*\*\**p* < 0.0001). **H.** Representative images of localization of Endonuclease G to mitochondrial matrix in HeLa cells. Conjugated secondary antibodies were used for detecting Endonuclease G (Alexa-488) and mitochondrial matrix protein, TFAM (Alexa-568). DAPI is used as nuclear stain. **I.** Colocalization analyses of Endonuclease G and TFAM signals using JaCoP in ImageJ software. Minimum of 50 cells were used for analysis of colocalization of red and green signals and plotted as the colocalization value of green overlapping red. Y-axis depicts the Mander’s colocalization coefficient value calculated for green over red and plotted in the form of dot plot. Control represents the panel where only one of the primary antibodies was used. **J.** Western blotting showing immunodepletion of Endonuclease G from rat testicular mitochondrial extracts. Beads were incubated with anti-Endonuclease G and then with the extracts. Actin served as a loading control. **K.** Immunodepletion of another endonuclease present in mitochondria, MGME1 from rat testicular mitochondrial extracts as described in panel J. **L.** Endonuclease G or MGME1- depleted extract was incubated with pDI1 and used for the primer extension using VKK11 primer. Extract without the addition of antibody served as bead control (lane 4). Lanes 3 is no protein control, lanes 5 and 6 corresponds to increasing concentrations of MGME1 depleted extracts, while in lanes 7 and 8, increasing concentrations of Endonuclease G depleted extracts were added. “M” is a 50 bp ladder. Cleavage positions are indicated by arrow and boxed. **M.** Reconstitution assay was performed by the addition of purified Endonuclease G following its immunodepletion. Lane 2 represents the primer alone; Lane 3 represents the beads control; lane 4 represents the Endonuclease G depleted extract while lane 5 and 6 represent the addition of purified Endonuclease G to the depleted extracts. **N.** Bar diagram representing the cleavage intensity after reconstitution assay. **O.** Knockdown of Endonuclease G from HeLa cell using PEI mediated transfection. shRNA against Endonuclease G cloned plasmid was used for transfection. Cells were harvested after 48 h and mitochondrial extracts were prepared. Western blotting was performed to confirm the knockdown of Endonuclease G from the HeLa cells. Actin served as a loading control. **P.** The knockdown extract was incubated with the plasmid and used for the primer extension using VKK11 primer (“I” and “II” are two biological repeats). Extract prepared from the sample transfected with scrambled plasmid served as a control. Lanes 3, 4, 7, 8 serve as scrambled controls while lanes 5, 6, 9, 10 are for knockdown extracts. “M” is a 50 bp ladder. **Q.** Bar diagram representing the cleavage intensity of the extracts prepared after transfection by scrambled plasmid and shEndo G plasmid. In panels N and Q, a minimum of three biological repeats were performed and the data is shown with the error bar calculated as SEM (ns: not significant, *p< 0.05, **p< 0.005, ***p< 0.0001).

For deciphering whether the position of cleavage observed was indeed at Region I, primer extension assay was carried on pDI1 and its mutant (pDI2) following cleavage by mitochondrial extracts isolated from testes and spleen. Results revealed two specific nicking sites when incubated with mitochondrial extracts from both testes and spleen, which were mapped to the position corresponding to the G-quadruplex forming region when DNA sequencing ladder was used as a marker (Figure 3C). Interestingly, mutation of the G4 motif resulted in a significant reduction of cleavage activity, as one of the two cleavage sites completely disappeared when 2 G stretches were mutated from G4 motif (Figure 3D). This may be due to G4 structure formation in alternate conformations, as Region I can fold into multiple confirmations (data not shown).

### Endonuclease G mediates structure-specific cleavage at mitochondrial Region I

Among different mitochondrial endonucleases, Endonuclease G was shown to bind preferentially to the guanine rich strands by one of the first studies (Ruiz-Carrillo and Renaud, 1987). Since we have observed the formation of G-quadruplex DNA structures at Region I of the mitochondrial genome, we considered the possibility of Endonuclease G as the protein responsible for the observed cleavage. In a different study, specific cleavage at kink DNA by Endonuclease G has also been reported (Ohsato et al., 2002). Based on these, we wondered whether Endonuclease G could cleave G4 DNA present at Region I on pDI1. To do this, Endonuclease G was overexpressed in bacteria, purified and identity was confirmed by western blotting (Figure S3A, B). Purified Endonuclease G was subjected to plasmid nicking assay and analysed on an agarose gel. Results showed that while the wild type plasmid (pDI1) showed a conformation change from supercoiled to open circular, mutant plasmid (pDI2) retained its supercoiled form (Figure 3E). Interestingly, primer extension studies revealed that Endonuclease G cleaved mitochondrial DNA at multiple sites corresponding to region of G- quadruplex DNA formation at Region I, when it was present on pDI1 (Figure 3F, lanes 1-3, 5-8). Importantly, the cleavage positions were comparable to the one observed, when mitochondrial extract was used for the study (Figures 3F, lane 4). More importantly, a mutation at the G- quadruplex forming motif abrogated cleavage by Endonuclease G (Figures 3D, S3C). Further, when endonucleases such as CtIP, FEN1 and RAGs were used for cleavage assay, results showed that unlike Endonuclease G, none of the purified endonucleases exhibited any significant cleavage (Figure S3J) although the presence of CtIP (Tadi et al., 2016) and FEN1 (Kalifa et al., 2009; Klungland and Lindahl, 1997; Tadi et al., 2016) has been shown previously in mitochondria. In all cases the identity and activity of the endonucleases used were confirmed prior to use (Figure S3D-I).

Considering that Endonuclease G is one of the important mitochondrial nucleases (Gole et al., 2015; McDermott-Roe et al., 2011; Misic et al., 2016; Wiehe et al., 2018; Zhou et al., 2016), its presence was confirmed in mitochondria, using mitochondrial specific marker, ‘MitoTracker Deep Red or MitoTracker green FM’ and using immunofluorescence with Endonuclease G antibody in three different cell lines of different origin (Figure 3G; S4A). Merge of MitoTracker with tagged Endonuclease G revealed localization of Endonuclease G in mitochondria in HeLa (human cervical cancer), HEK 293T (human embryonic kidney epithelial) and MEF (mouse embryonic fibroblast) cell lines (Figure 3G; S4A, B). Further, to check if Endonuclease G is present in the mitochondrial matrix, colocalization analysis was performed along with TFAM, a transcription factor known to be present in the mitochondrial matrix and Endonuclease G using immunofluorescence in HeLa cells (Figure 3H). Results revealed low but distinct levels of colocalization of Endonuclease G and TFAM, suggesting the presence of Endonuclease G in the mitochondrial matrix (Figure 3H, I). This was further confirmed by western blotting after fractionating the mitochondria (see below).

Immunodepletion of Endonuclease G in mitochondrial extracts prepared from rat testes showed a significant reduction in cleavage efficiency at the mitochondrial G-quadruplex Region I, compared to beads alone control and another important nuclease MGME1, known to be present in mitochondria (Figure 3J-L, S4C) (Phillips et al., 2017; Yang et al., 2018). Importantly, the activity was restored when purified Endonuclease G was added back following its immunodepletion from the extract, confirming the role of Endonuclease G in cleavage (Figure 3M, N). Further, shRNA mediated knockdown of Endonuclease G was performed in HeLa cells. Knockdown efficiency was assessed by western blotting using Actin as a loading control (Figure 3O). HeLa cells transfected with scrambled plasmid served as transfection control. Cleavage assay using mitochondrial extracts following knockdown of Endonuclease G revealed a significant reduction in cleavage efficiency at Region I, as that of scrambled control suggesting its role in mitochondria in the context of cells (Figure 3P, Q).

### Endonuclease G binds to G-quadruplex forming regions of mitochondria inside cells

To investigate if Endonuclease G binds to G-quadruplex forming regions of mitochondria, G-quadruplex specific antibody, BG4 was used for colocalization studies along with Endonuclease G (Figures 4A, B; S5A). Yellow foci in the merged image suggested colocalization of BG4 to Endonuclease G. ∼100 cells were analysed for colocalization of anti-G4 and anti-Endonuclease G using the Mander’s colocalization coefficient, for red over green, was ∼0.32, while green over red was ∼0.21 suggesting the binding of Endonuclease G to G-quadruplex DNA structure (Figure S5A).

**Figure 4.**
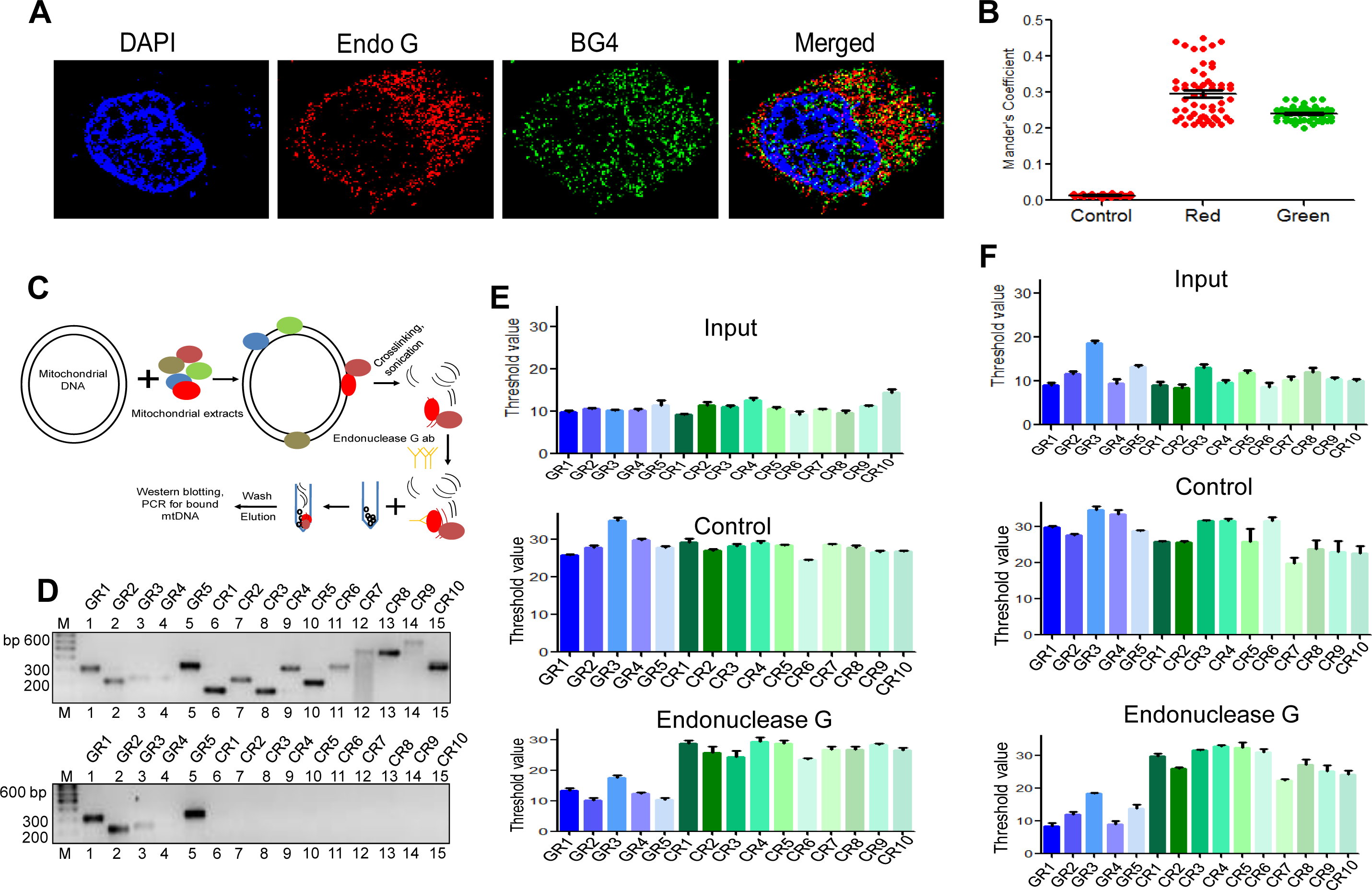
Investigation of binding efficacy of Endonuclease G to G4 DNA at Region I of mitochondrial genome. **A.** Representative image showing colocalization of Endonuclease G with BG4 in HeLa cells. Alexa-568 and Alexa-488 conjugated secondary antibodies were used for detection of Endonuclease G and BG4 proteins, respectively. DAPI was used as nuclear stain. **B.** The quantitation showing colocalization of Endonuclease G and BG4. The colocalization was quantified using Mander’s colocalization coefficient (ImageJ software) by analyzing a minimum of 100 cells and presented as a dot plot. Red plot represents the overlapping of Endonuclease G over BG4 while Green plot represents the overlapping of BG4 over Endonuclease G. **C.** Schematic showing the ChIP assay used for binding of Endonuclease G present in the rat testicular mitochondrial extracts to the mitochondrial genome. Bound regions were pulled out using anti-Endonuclease G and protein A/G beads. Regions of interest were detected by either semi-quantitative PCR or real-time PCR using appropriate primers. **D.** Agarose gel profile showing the amplification through semi-quantitative PCR of Input DNA (upper panel) and Endonuclease G pull down DNA (lower panel). Primers specific to 5 G-quadruplex forming regions (GR1-GR5) and 10 random regions (CR1-CR10) were also used for the amplification. **E.** Real-time PCR of 5 G-quadruplex forming regions (blue) and 10 random regions (green) following ChIP. Input DNA served as template control. Antibody control served as a negative control. Error bar represents three independent biological repeats. **F.** Evaluation of binding of Endonuclease G to different regions of the mitochondrial genome within cells by ChIP. Cells were crosslinked and then mitochondria were isolated. Endonuclease G bound DNA was obtained and was amplified for different regions of mitochondria, as explained in panel E. Graph is plotted for the Ct values obtained following real-time PCR as described above. The error bar represents three independent biological repeats.

To examine the specific binding of Endonuclease G to mitochondrial G quadruplex regions, an antibody binding assay was performed. Purified Endonuclease G was allowed to bind to mitochondrial DNA isolated from Nalm6 cells (Figure S5B) and crosslinked. Sonicated mitochondrial DNA was incubated with anti-Endonuclease G and Protein A/G beads. Resulting bound DNA was reverse crosslinked, purified and used for both real-time and semiquantitative PCR (Figure S5C, D). The threshold values presented as a bar diagram suggested that Endonuclease G can bind to all G quadruplex regions (GR1- GR5), including the one present at mitochondrial Region 1, while the control regions (CR1-CR10) (not known to form G quadruplex) did not bind to Endonuclease G (Figure S5D-F). These results reveal the specific binding of Endonuclease G to mitochondrial G quadruplex regions.

Further, the mitochondrial extract was incubated with the mitochondrial DNA and pull- down assay was done using the Endonuclease G antibody (Figure 4C). Following pull down, beads were separated and loaded for protein analysis and western to confirm the pull down (Figure S5F). Remaining was used for DNA isolation followed by real time PCR and semiquantitative PCR. Results suggested the binding of Endonuclease G, specifically to G quadruplex regions (Figure 4C-E). Endonuclease G binding assay inside cells was performed to assess the binding of Endonuclease G to mitochondrial G4 DNA (Figure S2C, S5C). The antibody bound mitochondrial DNA was used for real time PCR to amplify the G quadruplex regions and control regions. Results showed specific amplification of G quadruplex regions and no amplification of control regions, suggesting the specificity of Endonuclease G to G quadruplex regions within the cells (Figure 4F).

P1 nuclease assay was also performed to determine the precise binding footprint of Endonuclease G to G-quadruplex forming Region I by incubating the wild type and the mutant oligomers of Region I in presence and absence of Endonuclease G. Interestingly, binding of Endonuclease G to G-rich strand (G1) rescued the sensitivity of P1 nuclease (Figure S5G, lanes 5-8), while in case of C-rich strand, mutants and random oligomers, such a protection was absent (Figure S5G).

### Occurrence of MMEJ can explain the deletions in mitochondrial Region I

MMEJ is an alternate form of classical NHEJ, where sequence containing microhomology/direct repeats are used for joining of the broken ends resulting in deletions (Baldacci et al., 2015; Krasich and Copeland, 2017; Sharma et al., 2015; Tadi et al., 2016; Wang et al., 2014; Wisnovsky et al., 2018). G-quadruplex motif present in mitochondrial Region I contain a 9-bp repeat (CCCCCTCTA), and the deletion of one of the repeats was characteristic of many mitochondrial disorders. Based on previous studies, suggesting MMEJ as a DSB repair mechanism responsible for the occurrence of deletions in the mitochondrial genome (Tadi et al., 2016), the potential involvement of MMEJ was tested at the 9-bp deletion site. For this oligomeric DNA containing 7 bp repeat from the mitochondrial region along with flanking DNA sequence was synthesized and MMEJ assay was performed by incubating with mitochondrial extract from rat testes and analyzed (Figure 5A). If the joining occurred by utilizing 7 bp microhomology regions, it would give rise to a 105 bp fragment upon PCR amplification. Results showed that mitochondrial extracts catalysed joining at Region I by using microhomology (Figure 5B). Interestingly, joined products of lower sizes were also observed. Cloning and sequencing of PCR products revealed that all the clones analysed had deletions involving the microhomology region of 7 bp along with the flanking sequence suggesting the use of MMEJ mediated repair (Figure 5C). These results show that, indeed MMEJ operates when a DSB is generated at mitochondrial Region I and can explain the shorter mitochondrial deletions.

**Figure 5.**
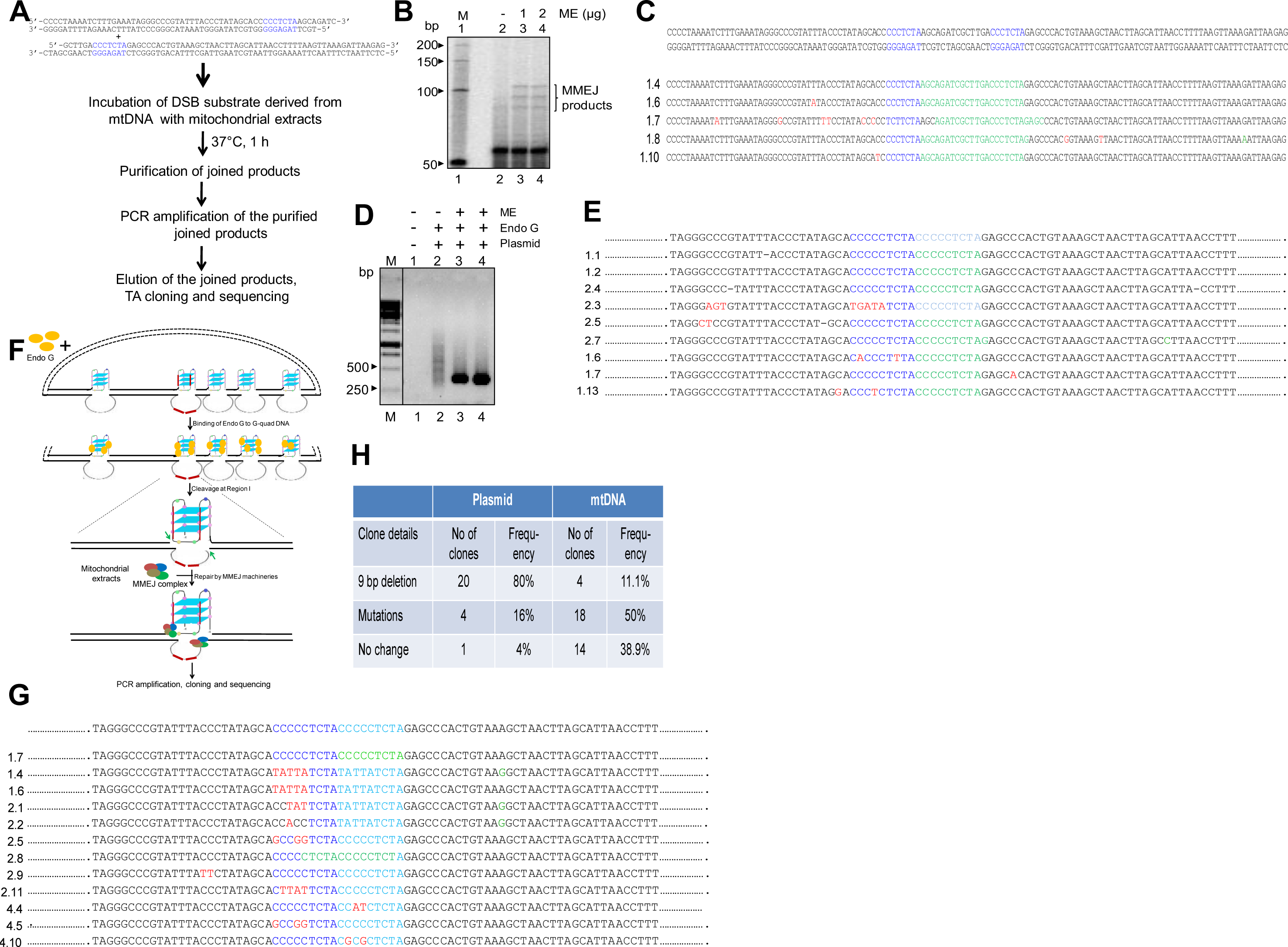
Assessment of involvement of DSB repair pathways during “9-bp deletion” associated with mitochondrial genome and its reconstitution. **A.** Schematic showing the investigation of DSB repair mechanism associated with fragility of Region I of mitochondria. Complementary oligomers were annealed to generate dsDNA harboring 7 bp direct repeats derived from Region I with overhang and was incubated with testicular mitochondrial extracts. The joined products were eluted and cloned into TA vector following PCR and sequenced to investigate the joining mechanism. **B.** Gel profile (8% denaturing PAGE) of the joined products after PCR amplification (using radiolabelled primer). The gel purified product was used for cloning and sequencing. **C.** Representative junctional sequences obtained after cloning and sequencing of EJ junctions. Blue sequences denote the direct repeats. Deleted sequences are denoted in green, while the mutated sequences are marked in red. **D**. Agarose gel profile of the joined product after PCR amplification of end joined DNA. Briefly, the plasmid DNA was incubated with Endonuclease G and the linearized pDNA was purified and the joining reaction was performed in the presence of mitochondrial extracts. The reaction without extracts served as control. The PCR products were then gel purified, cloned and sequenced. **E.** Representative sequences of end joined junctions following DNA sequencing of clones described in panel D. The direct repeats are in blue. Deleted sequences are in green and the mutated sequences are in red. **F.** Strategy used for reconstitution of ‘9-bp deletion’ associated with mitochondrial genome using purified Endonuclease G. Briefly, the mitochondrial DNA was incubated with purified Endonuclease G and the DNA was purified and precipitated. The linearized mtDNA was used for the joining reaction in the presence of mitochondrial extracts. The joined product was subjected to PCR using appropriate primers to amplify Region I, then cloned and sequenced as described above. **G.** Representative sequences of joining junction following reconstitution of “9- bp deletion”. For other details refer panel C and E. **H.** Table showing the frequency of clone obtained with ‘9-bp deletion’ or other mutations, when either plasmid or mtDNA was treated with Endonuclease G followed by repair catalyzed by mitochondrial extracts.

### Reconstitution of 9-bp deletion in the mitochondrial genome

Studies thus far revealed the involvement of Endonuclease G by binding and introducing cut at or near G4 DNA present at Region I of the mitochondrial genome. The breaks introduced by Endonuclease G are repaired using MMEJ, explaining generation of the deletion. In order to reconstitute the whole process of deletion formation in the mitochondrial genome, we have used two approaches. Either plasmid DNA containing mitochondrial Region I or purified mitochondrial genome were treated with purified Endonuclease G to generate DSBs (Figures S6A; 5F). The broken DNA was then purified, subjected to MMEJ mediated repair using mitochondrial extract from testes and further analysed. Interestingly, the deletion of 9-bp direct repeats from Region I was observed in 20 out of 25 clones sequenced, suggesting the utilization of the microhomology for repair (Figure 5D, E). Interestingly, there were mutations in the G-quadruplex forming motif in those clones where microhomology mediated joining was not seen (Figure 5E, clone number 2.3). Besides, when a similar strategy was used for mitochondrial DNA, 4 out of 37 clones had the 9-bp direct repeat deletion (Figures 5G, H; S6B). In these cases, few clones had mutation in G quadruplex region, while many others had sequences exactly like the mitochondrial genome.

Based on the reconstitution assay and previous studies (García-Lepe and Bermúdez-Cruz, 2019; Tadi et al., 2016; Wisnovsky et al., 2018), we propose that following induction of DNA breaks by Endonuclease G at Region I of the mitochondrial genome, DNA modifying enzymes such as CtIP, MRE11, Exonuclease G may help in generating 3’ overhangs. The microhomology regions are then aligned, and the gap is filled by Polθ following which ligation of broken ends by Ligase III/XRCC1 leading to the ‘9-bp deletion’ at the Region I of the mitochondrial genome (Figure 6). Although the expression of all proteins associated with MMEJ are not well established, it is likely that these proteins are responsible for alt-NHEJ in mitochondria. Consistent to this, significant upregulation of genes associated with MMEJ was observed in hepatocellular carcinoma (HCC), where ‘9-bp deletion’ has been reported (Figure S7A, B). Further, 271 out of 371 HCC patients showed upregulation of Endonuclease G in this case (Figure S7A, B).

**Figure 6.**
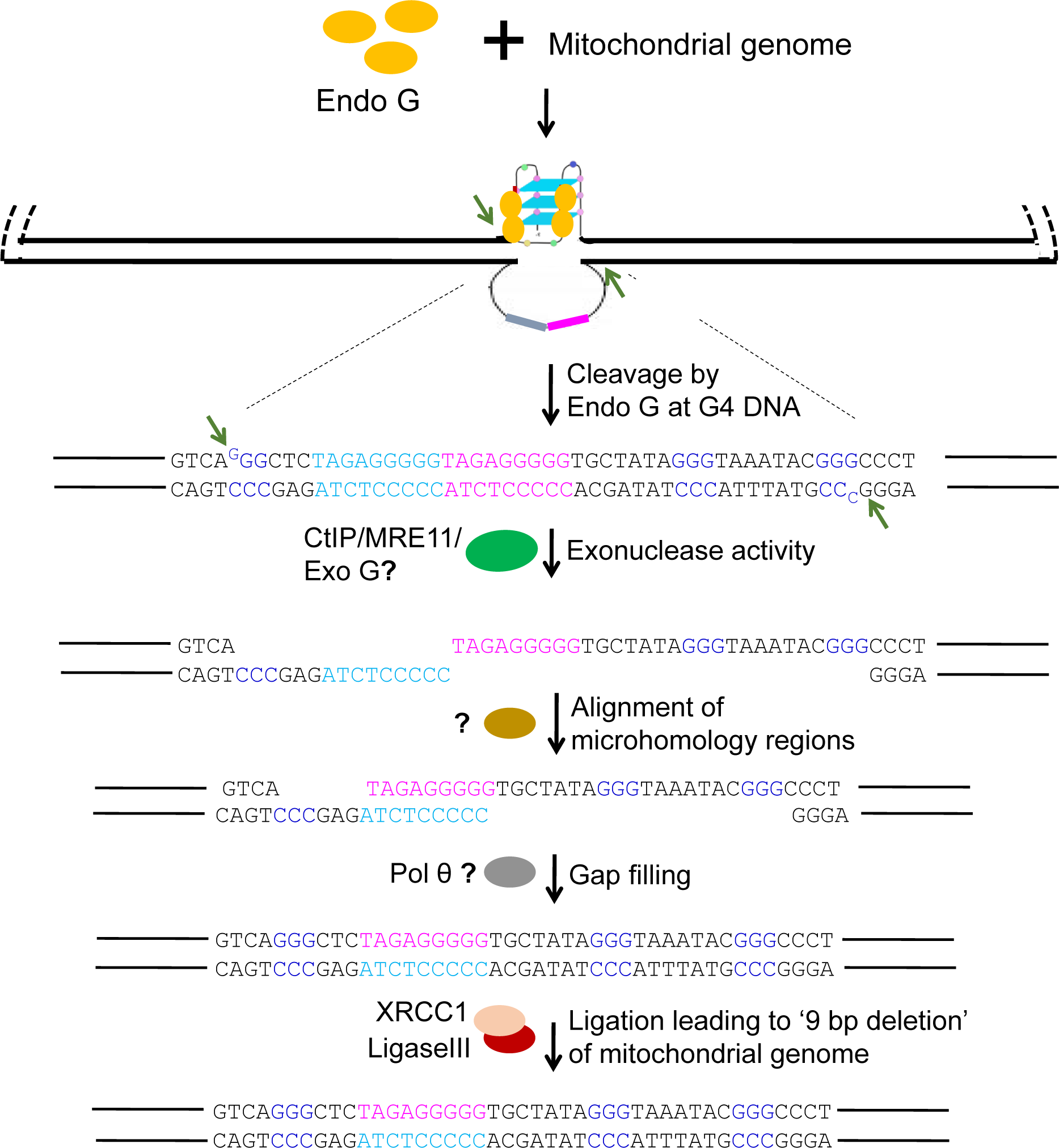
Model depicting mechanism of generation of ‘9-bp deletion’ seen in the mitochondrial genome. Endonuclease G binds and induces cleavage at single-double stranded junctions of G4 DNA as indicated by arrows. Exonuclease action (CtIP/MRE11/Exo G) exposes the direct repeats/microhomology region, which is then paired with the help of unknown proteins. Polθ mediated gap-filling followed by ligation of broken ends by Ligase III/XRCC1 can result in a ‘9-bp deletion’ at the Region I of the mitochondrial genome.

### Conditions that induce stress in mitochondria can facilitate release of Endonuclease G to mitochondrial matrix

Although Endonuclease G was thought to be localized within the mitochondrial intermembrane space, its proximity to the mitochondrial matrix was supported by multiple experimental evidence suggesting direct interaction of Endonuclease G with mtDNA (Duguay and Smiley, 2013; McDermott-Roe et al., 2011; Wiehe et al., 2018). We observed low level of colocalization of Endonuclease G with mitochondrial matrix protein, TFAM (Figure 3H, I). In order to examine this further we fractionated the inner and matrix proteins from the mitochondria by high speed ultracentrifugation. Western blotting analysis showed low, but distinct level of Endonuclease G in the mitochondrial matrix (Figure 7A).

**Figure 7.**
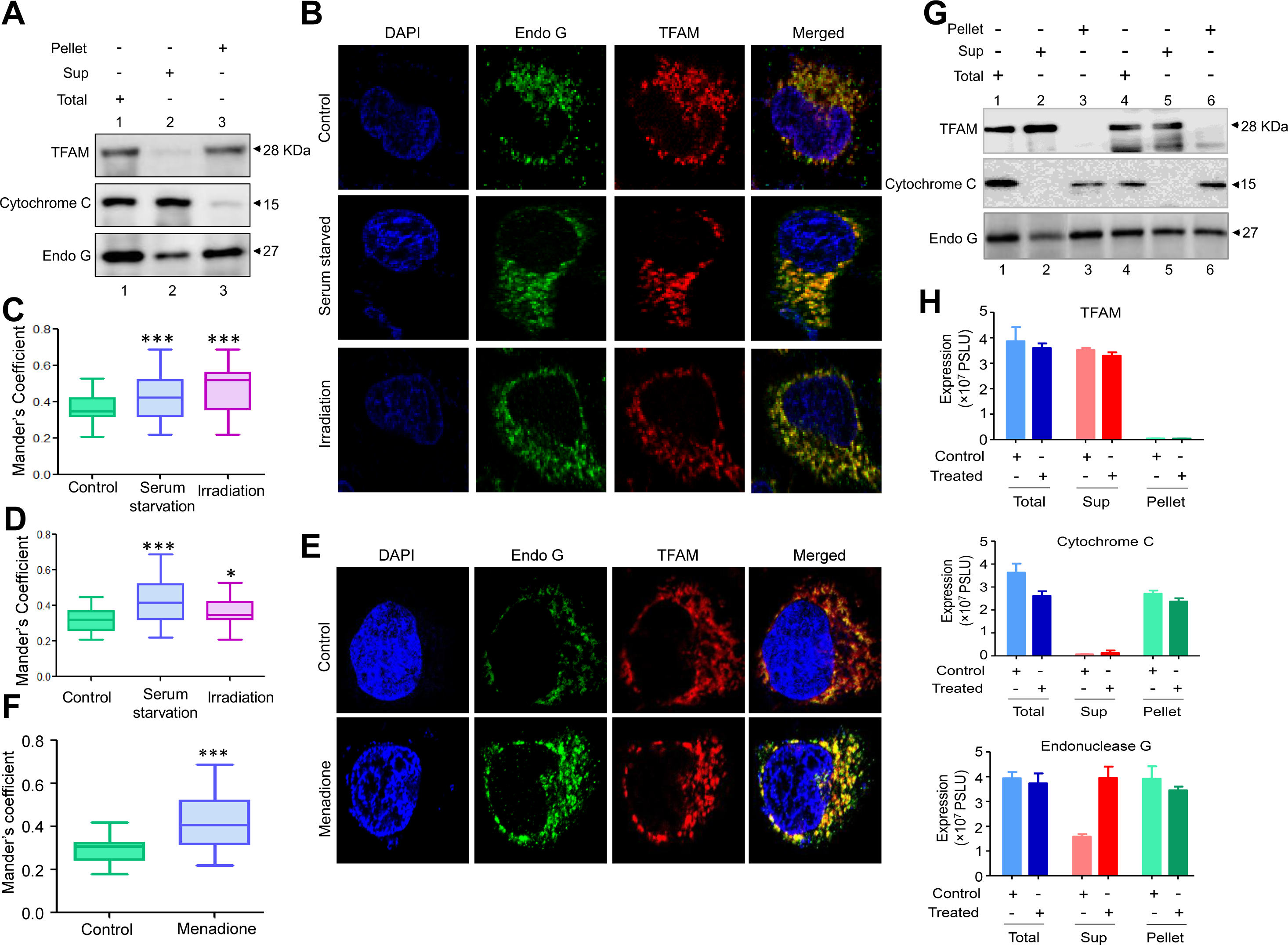
Investigation of stress conditions that favor the transport of Endonuclease G to mitochondrial matrix. **A**. Western blot showing the presence of Endonuclease G, Cytochrome C and TFAM in either extracts of total mitochondria, supernatant (Sup) and pellet. Sub- fractionation was performed using the standard high salt and sonication method followed by Ultracentrifugation. **B.** Representative immunofluorescence colocalization image of Endonuclease G with TFAM in HeLa cells under different stress conditions (serum starvation and Irradiation). **C**. Quantitation showing colocalization of Endonuclease G with TFAM in HeLa cells under different stress conditions. Data is presented as Box and Whiskers plot. **D.** Box and Whiskers plot representing quantitation of colocalization of Endonuclease G with TFAM in HCT cells under different stress conditions. **E**. Representative immunofluorescence colocalization image of Endonuclease G with TFAM in HeLa cells after treatment with menadione to induce mitochondrial stress. **F**. Quantitation showing colocalization of Endonuclease G with TFAM in HeLa cells when treated with Menadione. Data is presented as Box and Whiskers plot. **G**. Western blot showing the presence of Endonuclease G, Cytochrome C and TFAM in either total or supernatant (Sup) and pellet fraction with or without menadione treatment (25 µM) following sub-fractionation of mitochondrial compartments. Lanes 1-3 are for control samples and Lanes 4-6 are menadione treated samples. **H**. Bar graph showing the quantitation of presence of Endonuclease G, Cytochrome C and TFAM in either total mitochondria or supernatant (Sup) and pellet fraction with or without menadione treatment (25 µM). Quantitation is based on three biological repeats and the data is shown with the error bar calculated as SEM.

Next, we were interested in testing the conditions in which Endonuclease G is released to the mitochondrial matrix from the inner mitochondrial membrane. To do this, we exposed 2 different cell lines, HeLa and HCT116, to different stress conditions such as exposure to low dose irradiation (2 Gy) and serum starvation for 24 h. Cells were harvested and subjected to immunofluorescence analysis (Figure 7B). Results showed significant enhancement in the level of Endonuclease G in the matrix following exposure to IR (1.7 fold) and serum starvation (1.2 fold) compared to that of controls upon colocalization analysis in HeLa cells (Figure 7B, C). This was 1.3 fold and 1.4 fold, respectively in HCT116 cells (Figure 7D). Further, menadione, a mitochondria specific ROS inducer was used for the study in HeLa cells. Results showed 1.4 fold increase in Endonuclease G levels in mitochondrial matrix when cells were exposed to menadione (25 µM) for 2 h (Figure 7E, F). These results indicate that release of Endonuclease G can be regulated by stress conditions, which in turn could influence the incidence of mitochondrial deletions.

To further test above results, fractionation of mitochondria was performed as described above following exposure to IR (2 Gy) and Menadione (25 µM). Fractionated extracts were equalized and used for western blotting analysis (Figure 7G). Results showed that exposure to stress conditions resulted in elevated levels of Endonuclease G in the matrix and thus proximity to the mitochondrial genome (Figure 7G, H). It is possible this may act as a regulatory step in the generation of mitochondrial deletion. However, this warrants further investigation.

## DISCUSSION

In the present study, we have established existence of five G-quadruplex DNA structures in the mitochondrial genome, all with 3-plate conformation that follows the general empirical formula for G-quadruplexes d(G_3+_N_1-7_G_3+_N_1-7_G_3+_N_1-7_G_3+_) which is considered to be energetically more stable (Puig Lombardi and Londono-Vallejo, 2020). A close analysis of patient breakpoint regions revealed the presence of high-frequency mitochondrial deletion junctions (9 bp deletion) adjacent to one of the G-quadruplex structures at Region I.

Presence of G-quadruplex motifs proximal to mitochondrial deletions have been reported (Falabella et al., 2019; Oliveira et al., 2013). Computational analysis (Bharti et al., 2014) revealed that G4 sequences proximal to mtDNA deletions in a number of genetic diseases like Kearns-Sayre syndrome, Pearson marrow-pancreas syndrome, Mitochondrial myopathy, Progressive external ophthalmoplegia, etc. G4 motifs were also located adjacent to mitochondrial deletion breakpoints associated with KSS, a clinical subgroup of mitochondrial encephalomyopathies associated with these diseases (Van Goethem et al., 2003; Zeviani et al., 1988). In the present study, we establish that the region containing “9-bp deletion” can fold into G-quadruplex structures, as shown by EMSA and CD studies, which was K^+^ dependent. Using bisulphite modification assay, we showed that Region I can fold into a G4 DNA in the mitochondrial genome. IF studies and antibody pull down assays using G4 specific antibody, BG4 provide further evidence for the occurrence of G4 quadruplex DNA structures within the cells. Thus, with the combinatorial use of *in silico* studies, biophysical, biochemical and *ex vivo* techniques, we demonstrate the formation of G quadruplexes at the region corresponding to 9- bp deletion in mitochondrial genome.

Formation of intramolecular parallel G-quadruplex structure in mitochondrial genome could make the region fragile as there is an increased probability of replication fork slippage or transcription arrest at this region. Primer extension assay revealed multiple pause sites around G4 motif, suggesting that polymerase arrest could occur at these structures. However, the deletions seen were very specific and occurred in a particular sequence. Thus, it is highly unlikely that replication or transcription arrest contributes to a double-strand break and thus deletion. Thus, an alternate possibility for the generation of a break at these structures could be due to recognition by structure-specific nucleases followed by cleavage at G4 DNA.

Studies have shown that a number of enzymes possess the ability to cleave G-quadruplex structures. The action of RAGs at the G-quadruplex site in BCL2 MBR (Raghavan et al., 2004), MRE11 and DNA2 helicase on either side of the G-quadruplex structure *in vitro* (Masuda-Sasa et al., 2008) being some of the examples. In the current study, we observed that the structure-specific nuclease activity of mitochondrial endonuclease G can induce cleavage at G4 DNA formed at Region I of the mitochondrial genome using both biochemical (primer extension with purified nucleases, m-ChIP with purified Endonuclease G and isolated mitochondria, reconstitution assay with plasmid and purified protein) and *ex vivo* (immunofluorescence, mitoChIP, reconstitution assay in mitochondria) assays. Previous studies have reported various other properties of Endonuclease G, which include generation of primers during mitochondrial DNA replication by polymerase γ, initiation of genomic inversion in herpes simplex type-1 virus (HSV-1), and its role during apoptosis (Cote et al., 1989; Parrish et al., 2001; Ruiz-Carrillo and Renaud, 1987).

Mammalian Endonuclease G is encoded in the nucleus (Gannavaram et al., 2008; Ohsato et al., 2002; Wiehe et al., 2018; Zhou et al., 2016) and upon translocation to mitochondria, the mitochondrial targeting sequence is cleaved off such that a mature nuclease is released (Li et al., 2001; van Loo et al., 2001; Wiehe et al., 2018). Generally, Endonuclease G is known to reside at the intermembrane space of mitochondria, although a low expression has been reported in the matrix (Duguay and Smiley, 2013; McDermott-Roe et al., 2011; Wiehe et al., 2018). In the present study, we have observed a low level of Endonuclease G in the mitochondrial matrix. However, it is likely that Endonuclease G is further released to matrix when cells are under stress, or when exposed to ROS. In fact, our results reveal that upon menadione treatment, serum starvation or irradiation, level of Endonuclease G in the mitochondrial matrix was significantly high.

In a recent study, it has been shown that Endonuclease G also cleaved mtDNA in the case of unrepaired oxidative lesions (Wiehe et al., 2018). This was helpful in compensatory mtDNA replication through TWINKLE and mtSSB (Wiehe et al., 2018). However, binding of Endonuclease G to mtDNA and cleavage might be an event dependent upon various factors, including oxidative stress. Therefore, it is possible that in a normal cellular condition, level of Endonuclease G in the matrix is low, however upon stress, it is released into the matrix leading to genome fragility. Hence, stress-induced release may act as a regulatory mechanism with respect to the generation of mitochondrial deletions. A recent study showed the complete degradation property of Endonuclease G at pH 6 which could delineate its role during apoptosis, since apoptotic cells show cytosolic acidification. Interestingly, at physiological pH, the enzyme showed the property of nuclease (converting supercoiled DNA to open circular and linear) (Schafer et al., 2004). CPS-6, a mitochondrial endonuclease G in *Caenorhabditis elegans* acts with maternal autophagy and proteasome machineries to promote paternal mitochondrial elimination. It relocates from the intermembrane space of paternal mitochondria to the matrix after fertilization to degrade mitochondrial DNA (Zhou et al., 2016). *Endog^-/-^* mice revealed that Endonuclease G can act as a novel determinant of maladaptive cardiac hypertrophy, which is associated with mitochondrial dysfunction and depletion (McDermott-Roe et al., 2011).

Our study suggests that Endonuclease G can bind to G-quadruplexes and nick the DNA resulting in single-strand breaks, which can get converted as double-strand breaks during replication or during other DNA transactions. We observed that the binding was highly specific to G4 DNA, as we did not observe nonspecific binding to regions even when the regions where GC content was high in the mitochondrial genome. Previous studies have shown that although Endonuclease G showed random nuclease activity at higher concentrations, at optimal concentrations, it was specific to kink DNA (Ohsato et al., 2002). The first study on Endonuclease G had shown that it has preference towards the guanine rich strands, which falls in line with our study as the formation of G quadruplexes requires a G-rich region (Ruiz-Carrillo and Renaud, 1987). Further, Endonuclease G also cleaved mtDNA in the case of unrepaired oxidative lesions (Wiehe et al., 2018). It is likely that, binding and cleavage by Endonuclease G on mtDNA be an event dependent on multiple factors, including oxidative stress. Based on our data, it is evident that in the presence of stress, levels of Endonuclease G go up in the matrix, which enables the protein to be present proximal to the mitochondrial genome. The findings from m-ChIP studies suggest that Endonuclease G only binds to the G-quadruplex forming regions and not to other regions of mtDNA. This provides an additional regulation and ensure the stability of mitochondrial genome in most instances.

DNA repair in mitochondria is poorly understood. Base excision repair (BER) is the most well characterized DNA repair pathway in mitochondria, while it is generally believed that nucleotide excision repair is absent (Akbari et al., 2008; de Souza-Pinto et al., 2009; Liu et al., 2008). Homologous recombination repair helps in the repair of DSBs and thus stability of the genome (Dahal et al., 2018), though classical NHEJ was undetectable (Tadi et al., 2016). In contrast, microhomology mediated end joining uses flanking direct repeats to fix DSBs and sequence in between the repeats are deleted (Tadi et al., 2016). Interestingly, the G quadruplex forming Region I is present close to the deletion junction 8271:8281 (Redd et al., 1995; Yao et al., 2000) had also a 9 bp direct repeat sequence at the flank site. Our results demonstrate that that the break generated by Endonuclease G at G4 DNA in mitochondrial Region I is repaired through MMEJ and the direct repeats are used for joining followed by exonuclease action leading to deletion of in between DNA sequences. However, we observed that some of the clones had long mutated stretch, which could be due to failed long patch base excision repair (Szczesny et al., 2008).

In a recent study, a replication slippage mediated mechanism was attributed for a common 4977 deletion where in frequent replication fork stalling at the junction of the common deletion was observed (Phillips et al., 2017). While this is a replication dependent process, what we observed may not be immediately connected to replication. However, it is possible that replication can indeed facilitate the formation of G-quadruplexes.

A balance between the formation and resolution of G4 DNA within the mitochondrial genome is important. The mitochondrial genes are transcribed as a single polycistronic mRNA (Clayton, 1984; Sanchez et al., 2011) and therefore undergo a high rate of transcription, which may promote G-quadruplex formation. Besides, random ‘breathing’ of DNA, (a process of transient melting of the duplex structure due to thermal fluctuations) may also occur in the GC rich sequences of the mitochondrial DNA promoting G-quadruplex formation (Dornberger et al., 1999; Jose et al., 2009). Once the structure is formed, G4 resolvases also play a critical role in its resolution. In the nuclear DNA context, resolvases like WRN and BLM helicase are considered as the factors that negatively regulate G4 DNA (Giri et al., 2011; Paeschke et al., 2013). However, such a role for these nucleases in mitochondria is not reported. TWINKLE helicase is known to unwind mitochondrial duplex DNA and regulate the replication, although its role in structural resolution is yet to be studied (Phillips et al., 2017).

Levels of Endonuclease G could be another factor that may regulate the process of deletion. Considering that Endonuclease G level is normally very low in the matrix, it is possible that the generation of deletions may have a connection with stress induced within mitochondria at a given time as Endonuclease G is transported to the matrix at that time. If the deletion formation would have determined by replication, one would have anticipated such deletion to be seen in every molecule, which is not the case. Finally, the efficiency of MMEJ and factors that regulate transport of MMEJ proteins from nuclear to mitochondria also may play an important role.

Hence, our study suggests a novel mechanism that explains fragility in mitochondria, which is dependent on structure-specific nuclease activity of Endonuclease G on G-quadruplex DNA and repair through an error-prone repair of MMEJ. Although we have focused on a single example of ‘9-bp deletion’ often seen in mitochondria, it is very much possible that it may be a general mechanism and may help in explaining several other deletions seen in mitochondria- associated human disorders.

## EXPERIMENTAL PROCEDURES

### Enzymes, chemicals, and reagents

Chemicals and reagents used in the present study were purchased from Sigma Chemical Co. (USA), SRL (India), Himedia (India) and Amresco (USA). Restriction enzymes and other DNA-modifying enzymes were obtained from New England Biolabs (USA). Culture media was from Sera Laboratory International Limited (UK), Lonza (UK). Fetal bovine serum and PenStrep were from Gibco BRL (USA). Radioisotope-labelled nucleotides were from BRIT (India). Antibodies were purchased from Abcam (UK), Santa Cruz Biotechnology (USA), BD (USA), Cell Signaling Technology (USA) and Calbiochem (USA). Oligomeric DNA was purchased from Juniper Life Sciences (India) and Medauxin (India).

### Oligomeric DNA

The oligomers used in this study are listed in Table 1 and as a separate supplementary excel file. Oligomeric DNA were purified on 12-15% denaturing PAGE, when needed (Raghavan et al., 2005b; Tadi et al., 2016). The details of oligomers used for different experiments are provided in appropriate sections. A more simplified nomenclature is used for certain oligomers while explaining the results of some experiments.

**Table 1.**
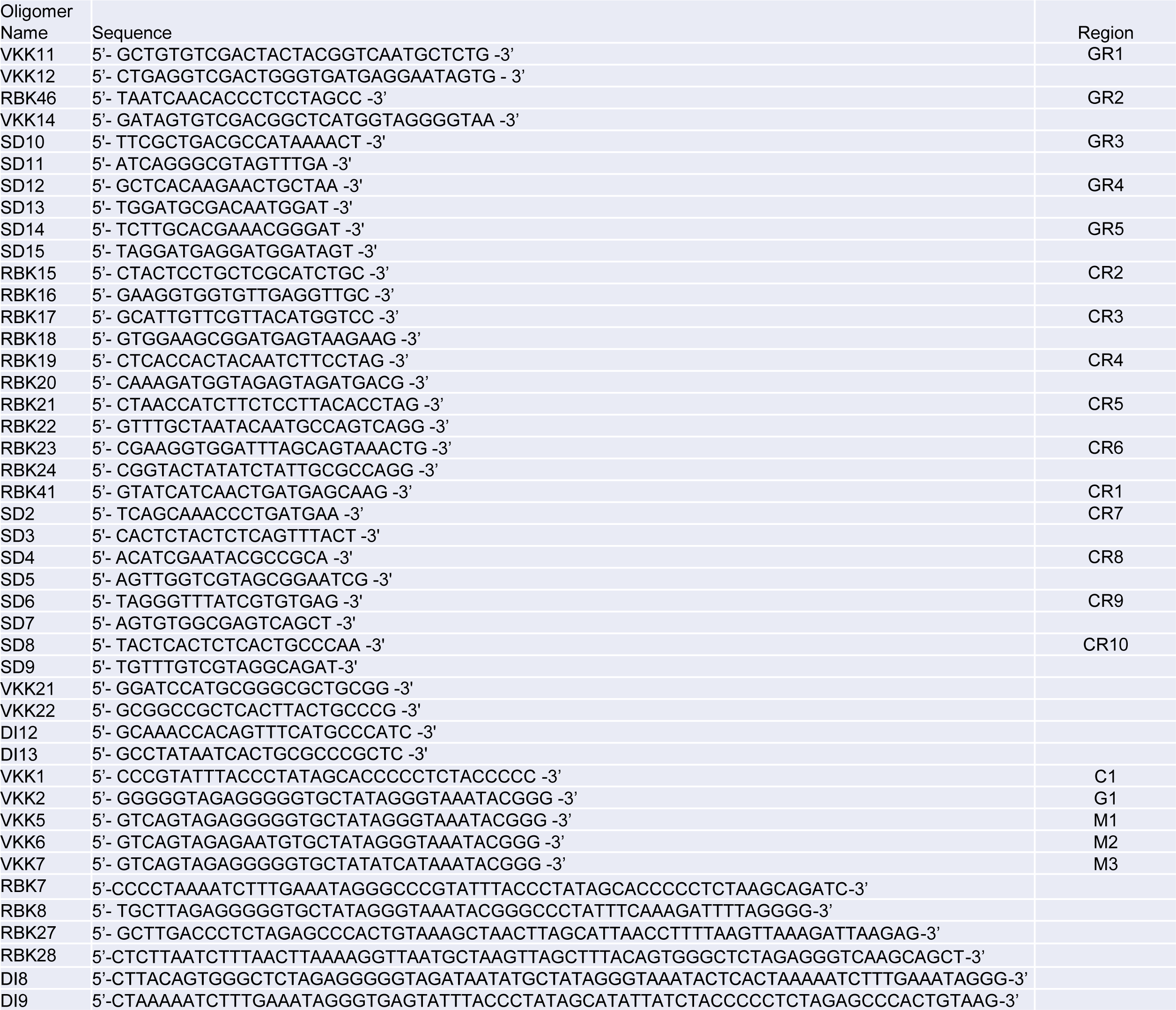
Oligomers used in the present study.

### Preparation of oligonucleotide dsDNA substrates

Oligomers RBK7 were annealed to RBK8 and RBK27 to RBK28 by incubating in the presence of 100 mM NaCl and 1 mM EDTA as described before (Kumar et al., 2010).

### Cell lines

HeLa (human cervical cancer), MEF (mouse embryonic fibroblast) and HEK293T (human embryonic kidney epithelial cell line) were purchased from National Centre for Cell Science, Pune, India. Nalm6 and Reh cells were from Dr. M. R. Lieber (USA) and Rho(0) cells from J. Neuzil (Australia). Cells were cultured in RPMI1640 or MEM medium supplemented with 10-15% fetal bovine serum (FBS), 100 μg/ml Penicillin, and 100 μg/ml streptomycin and incubated at 37°C in a humidified atmosphere containing 5% CO2 as described before (Srivastava et al., 2012). Rho(0) cells of B16 cells origin were cultured in DMEM medium supplemented with 15% fetal bovine serum (FBS), 100 μg/ml Penicillin, 100 μg/ml streptomycin, 50 mg/ml Uridine and 1 mM Sodium pyruvate and incubated as described above (Dong et al., 2017; Tan et al., 2015).

### Animals

Male Wistar rats (*Rattus norvegicus*) 4-6 weeks old were purchased from Central animal facility, Indian Institute of Science (IISc), Bangalore, India and maintained as per the guidelines of the animal ethical committee in accordance with Indian National Law on animal care and use (CAF/Ethics/526/2016).

### shRNA

shRNA used in the study was purchased from Resource Center at Division of Biological Science, IISc, Bangalore (funded by DBT: BT/PR4982/AGR/36/718/2012) Indian Institute of Science, Bangalore (India).

### Plasmid constructs

#### Plasmid construction for mitochondrial G-quadruplex assays

Mitochondrial DNA isolated from Nalm6 cells was amplified by primers specific for Region I (VKK11 and VKK12). The PCR products were purified, blunted and ligated into the EcoRV site of pBlueScript SK(+) to obtain the wild type plasmid for mitochondrial Region I (pDI1). Mutant plasmid for mitochondrial Region I was generated by PCR mediated site directed mutagenesis as described before (Raghavan et al., 2005c) using the primers VKK11, DI8, VKK12 and DI9 followed by cloning into pBlueScript SK(+). The resulting plasmid, named as pDI2, had a mutation in two G stretches of the Region I sequence.

#### Construction of Endonuclease G containing vector for purification of the protein

Endonuclease G coding sequence was amplified using primers VKK21 and VKK22, which were designed with BamHI and NotI restriction sites, respectively at flanking regions. PCR product was gel purified and cloned into EcoRV site of pBS SK+ cloning vector. For the generation of pRBK2, coding DNA sequence was digested from cloning vector using BamHI and NotI enzymes and cloned into the same site (BamHI and NotI) of pET28a+.

#### Plasmid isolation and purification

For each plasmid, after transformation, *Escherichia coli* was cultured in 500 ml of Luria broth (HiMedia, USA) for 18 h at 37°C. Isolation of plasmid DNA was performed by standard alkaline lysis method. It was then purified and precipitated, as described previously (Sambrook, 1989). The pellet was dissolved in TE (pH 8.0).

#### Isolation of mitochondrial DNA

Mitochondria were isolated from Nalm6 cells. Cells were first homogenized in the buffer [70 mM sucrose, 200 mM mannitol, 1 mM EDTA, and 10 mM HEPES (pH 7.4), 0.5% BSA per g tissue] in a Dounce-type homogenizer (20-30 strokes). Homogenate was centrifuged (3000 rpm, 10 min at 4°C) to remove nuclei and cellular debris from the supernatant that contains the cytosol and mitochondria. The supernatant fraction was centrifuged at 12,000 rpm (30 min at 4°C) to pellet the mitochondria. The mitochondrial pellet was washed in suspension buffer [10 mM of Tris–HCl (pH 6.7), 0.15 mM of MgCl2, 0.25 mM of sucrose, 1 mM PMSF and 1 mM DTT] twice and the mitochondrial pellet was incubated in the buffer containing 75 mM NaCl, 50 mM EDTA (pH 8.0), 1% SDS and 0.5 mg/ml Proteinase K to remove denatured proteins. The resulting lysate was deproteinized by phenol-chloroform extraction. The mitochondrial DNA was precipitated with 3 M Sodium acetate (pH 5.2) and chilled 100% ethanol. Purity of the mitochondrial DNA preparation was evaluated by PCR amplification with nuclear and mitochondrial DNA specific primers.

#### Preparation of mitochondrial extracts

Mitochondrial protein extracts were prepared following the isolation of mitochondria based on differential centrifugation as described (Dahal et al., 2018; Tadi et al., 2016). Testes and spleen from 4- to 6-week-old male Wistar rats were minced on ice and lysed in mitochondrial lysis buffer [50 mM of Tris–HCl (pH 7.5), 100 mM of NaCl, 10 mM of MgCl2, 0.2% Triton X-100, 2 mM of EGTA, 2 mM of EDTA, 1 mM of DTT and 10% glycerol] by mixing (30 min at 4°C) along with protease inhibitors (PMSF (1 mM), aprotinin (1 μg/ml), pepstatin (1 μg/ml) and leupeptin (1 μg/ml). The mitochondrial extract was centrifuged at 12,000 rpm (5 min) and the supernatant (mitochondrial fraction) was aliquoted, snap-frozen and stored at −80°C till use.

Mitochondrial protein extracts were also prepared from rat tissues as per manufacturer’s instructions using mitochondrial extraction kit (Imgenex, USA). Purity of the mitochondrial extracts prepared was checked by immunoblotting using the nuclear and mitochondrial specific markers.

#### Fractionation of mitochondria

4 × 10^7^ HeLa cells were treated with 25 µM Menadione and incubated for 2 h at 37°C. The mitochondrial pellet was then subjected to fractionation in high salt conditions (500 mM NaCl and 10 mM Tris). Mitochondrial pellet was then sonicated for 10 min at 37% duty per cycle. The samples were then separated into membrane and matrix fractions by ultracentrifugation at 100 000 g (TLA 110 rotor) for 30 min at 4°C in OptimaTM TLX table-top ultracentrifuge (Beckman-Coulter). The supernatant fraction was transferred to new tube. To the pellet, 1X PBS was added and both the fractions were used for western blotting analysis (Sinha et al., 2010; Suzuki et al., 2002).

#### 5’ end labeling of oligomeric substrates

The 5’ end labelling of oligomeric substrates was done using γ[^32^P]ATP in the presence of 1 U of T4 Polynucleotide kinase in a buffer containing 20 mM Tris-acetate [pH 7.9], 10 mM magnesium acetate, 50 mM potassium acetate and 1 mM DTT at 37°C for 1 h. The radiolabelled oligomers were purified on a Sephadex G25 column and stored at -20°C until further use (Kumar et al., 2010; Tadi et al., 2016).

#### Gel mobility shift assays

Gel mobility shift assays were performed using 5’ radiolabelled oligomeric DNA substrates (4 nM) predicted to form G-quadruplex structures, its complementary oligos and the mutants by incubating in a buffer containing 10 mM Tris-HCl (pH 8.0), 1 mM EDTA, with or without 100 mM KCl at 37°C and resolved on 15% native polyacrylamide gels. In conditions where the effect of KCl was studied, 100 mM KCl was added while preparing gels and the running buffer (1X TBE). Electrophoresis was carried out at 150 V at RT. The dried gels were exposed and signals were detected by phosphorImager FLA9000 (Fuji, Japan).

For BG4 binding assay, 5’ radiolabelled oligomeric DNA substrates (4 nM) predicted to form G-quadruplex structures and its complementary oligomer was incubated in a buffer containing 10 mM Tris-HCl (pH 8.0), 1 mM EDTA, with 100 mM KCl at 37°C for 1 h (Das et al., 2016; Javadekar et al., 2020). It was then incubated along with increasing concentrations of purified BG4 protein (100, 200, 400 and 800 ng) in a buffer containing 250 mM Tris (pH 8.0), 1 mM EDTA, 0.5% Triton X-100, 500 μg/ml BSA and 20 mM DTT for 1 h at 4°C. The complex was then resolved on 5% native polyacrylamide gels in the presence of 100 mM KCl in gel and the running buffer (1X TBE). Electrophoresis was carried out at 100 V at 4°C. The dried gel was exposed to a PI cassette and the radioactive signals were detected by phosphorImager FLA9000 (Fuji, Japan).

#### Circular dichroism (CD)

Circular dichroism studies for analysis of mitochondrial G-quadruplex formation were performed with oligonucleotide substrates (2 µM) corresponding to G-rich sequences and complementary C-rich sequences in a buffer containing 10 mM Tris-HCl (pH 8.0) and 1 mM EDTA. The experiments were set up either in the presence or absence of 100 mM KCl. The spectra were recorded between wavelengths 200 and 300 nm (5 cycles, scan speed of 50 nm/sec, RT) using JASCO J-810 spectropolarimeter (Nambiar and Raghavan, 2012; Raghavan et al., 2005a), and analyzed using SpectraManager (JASCO J-810 spectropolarimeter).

#### DMS protection assay

The radiolabelled oligomer VKK19 was incubated with Dimethyl Sulphate (1:250 dilution) in buffer containing 10 mM Tris-HCl (pH 8.0) and 1 mM EDTA either in the presence or absence of 100 mM KCl, at RT for 15 min (Nambiar et al., 2013). An equal volume of 10% piperidine was added, and the reactions were incubated at 90°C for 30 min. The reaction mixtures were diluted 2-fold with double distilled water and vacuum dried in a Speed Vac concentrator. The resulting pellet was again washed with water and vacuum dried. This procedure was repeated thrice. The pellet finally resuspended in 10 μl TE buffer (10 mM Tris-HCl, (pH 8.0) and 1 mM EDTA) was mixed with formamide containing dye and resolved on a 15% denaturing PAGE.

#### Polymerase stop assays

Plasmid substrates pDI1, pDI2 (mitochondrial Region I), Vent Exo(-) polymerase was used for primer extension assays (Kumari et al., 2015; Nambiar et al., 2013) in a buffer containing 20 mM Tris-HCl (pH 8.8), 10 mM (NH4)2SO4,10 mM KCl, 2 mM MgSO4, 0.1% Triton X-100 with additional supplementation with 75 mM KCl, LiCl or NaCl where required. After addition of 200 μM dNTPs, 0.2 U of Vent Exo(-) and 5’ radiolabelled primers (VKK11, VKK12, DI12), the reaction was carried out in a one-step PCR mediated primer extension assay for mitochondrial Region I (95°C for 10 min, 55-65°C for 3 min as specified below for each primer and 75°C for 20 min, as a single cycle) or multi cycle PCR extension (Denaturation at 95°C for 5 min, followed by 95°C for 45 sec, 55°C for 45 sec and 72°C for 45 sec over 20 cycles and a final extension at 72°C for 5 min). The reaction products were then loaded on 8% denaturing PAGE and the products were visualized as described above.

#### Preparation of sequencing ladders

Sequencing ladders for the primers VKK11 and VKK12, were prepared using the cycle sequencing method with a dNTP:ddNTP ratio of 1:20, 1:40, 1:30 and 1:10 for C, T, A and G ladders, respectively. The dNTP:ddNTP mix were separately provided in reaction mixtures containing 10 nM plasmid template, 0.5 μM 5’ radiolabelled primers, 20 mM Tris-HCl (pH 8.8), 10 mM (NH4)2SO4, 10 mM KCl, 2 mM MgSO4, 0.1% Triton X-100 and 1 U of Vent Exo (-) polymerase. The PCR was carried out using the following conditions: Step 1 at 95°C for 30 sec, 60°C for 30 sec and 72°C for 1 min (25 cycles) followed by Step 2 at 95°C for 30 sec and 72°C for 2 min (10 cycles).

#### Sodium bisulphite modification assay

pDI1, plasmid containing mitochondrial Region I or mitochondrial DNA isolated from Nalm6 cells were treated with sodium bisulfite as described earlier (Raghavan et al., 2004; Raghavan et al., 2006). Briefly, approximately 2 μg of mitochondrial DNA was treated with 12.5 μl of 20 mM hydroquinone and 458 μl of 2.5 M sodium bisulfite (pH 5.2) at 37°C for 16 h. The bisulphite treated DNA was purified using Wizard DNA Clean-Up Kit (Promega, Madison, WI) and desulfonated by treating with 0.3 M NaOH (15 min at 37°C). The DNA was ethanol precipitated and resuspended in 20 μl TE buffer. The mitochondrial Region I was PCR amplified, resolved on 1% agarose gels, purified and TA cloned. Clones were sequenced to analyze for conversions. The experiment was repeated 3 independent times and the cumulative data is presented.

### Overexpression and purification of endonucleases

#### Endonuclease G

For expression of protein, *E. coli* Rosetta cells were transformed with pRBK2 and grown till OD reaches 0.6 and then induced with 1 mM IPTG at 37°C for 3 h. Cells were harvested and the extract was prepared using extraction buffer (20 mM Tris-HCl (pH 8.0), 0.5 M KCl, 20 mM imidazole (pH 7.0), 20 mM Mercaptoethanol, 10% glycerol, 0.2% Tween 20, 1 mM PMSF) by sonication followed by loading on to Ni-IDA column (Macherey-Nagel, Germany). Endonuclease G was eluted in gradient imidazole concentration (100-500 mM) and pure fractions were pooled and the identity of the protein was confirmed by immunoblotting and used for the assays.

#### RAGs

MBP cRAGs (RAG1, amino acids 384–1040; RAG2, amino acids 1–383) were purified using a method as described previously (Naik et al., 2010; Raghavan et al., 2005b). Briefly, 293T cells were transfected with 10 μg of plasmid by the calcium phosphate method. After 48 h of transfection, cells were harvested, and proteins were purified using amylose resin column (New England Biolabs). Fractions were eluted and checked by silver staining. The activity was checked by nicking assay on standard recombination signal sequence substrate (AKN1/2) (Nishana and Raghavan, 2012).

#### FEN1

FEN1 was purified using expression plasmid pET-FEN1 CH as described previously (Greene et al., 1999). Briefly, culture was grown till OD reaches ∼0.5, induced with 0.5 mM IPTG at 37°C for 3 h. Cells were harvested, lysed and the lysate was loaded on to Ni-NTA column. FEN1 was eluted in increasing imidazole concentrations (100-500 mM) and pure fractions were pooled, and identity of the protein was confirmed by immunoblotting and used for the assays.

#### CtIP

CtIP was purified using expression plasmid pET14b-CtIP as described previously (Yu and Baer, 2000). Briefly, culture was grown till OD reaches 0.6, induced with 0.5 mM IPTG at 37°C for 4 h. Cells were harvested, lysed and the lysate was loaded on to Ni-NTA column. FEN1 was eluted in increasing imidazole concentrations (100-800 mM) and pure fractions were pooled and the identity of the protein was confirmed by immunoblotting and used for the assays.

#### Overexpression and purification of BG4

The plasmid expressing BG4 protein, pSANG10-3F-BG4 was a gift from Shankar Balasubramanian (Addgene plasmid # 55756). The plasmid was transformed into *E. coli*, BL21 (DE3), and the culture was expanded by incubating at 30°C, till the O.D. reached upto 0.6 (Biffi et al., 2013; Das et al., 2016; Javadekar et al., 2020). The cells were then induced with 1 mM IPTG for a period of 16 h at 16^°^C, harvested, and resuspended in lysis buffer (20 mM Tris-HCl [pH 8.0], 50 mM NaCl, 5% glycerol, 1% Triton X-100 and 1 mM PMSF). The cells were lysed by sonication, centrifuged and the supernatant was then loaded onto a Ni-NTA chromatography column (Novagen, USA). BG4 was eluted using increasing concentrations of Imidazole (100- 400 mM). BG4 enriched fractions were dialyzed against dialysis buffer (PBS containing 0.05% Triton X-100, 1 mM Mercaptoethanol, 5% glycerol and 0.1 mM PMSF) overnight at 4°C. Identity of the protein was confirmed by immunoblotting, using an anti-FLAG antibody (Calbiochem, USA) (Kumari et al., 2019). Activity assay for each batch of BG4 was checked by performing binding assay as described previously (Javadekar et al., 2020).

#### Immunoblot analysis

For immunoblotting analysis, approximately 20-30 μg protein was resolved on 8-10% SDS-PAGE (Chiruvella et al., 2008; Kumar et al., 2010). Following electrophoresis, proteins were transferred to PVDF membrane (Millipore, USA), blocked using 5% non-fat milk or BSA in PBS with 0.1% Tween-20. Proteins were detected with appropriate primary antibodies against Cytochrome-C, PCNA, Tubulin, CtIP, FEN1, RAG, Endonuclease G and appropriate secondary antibodies as per standard protocol. The blots were developed using chemiluminescent substrate (Immobilon™ western, Millipore, USA) and scanned by gel documentation system (LAS 3000, FUJI, Japan).

#### Immunoprecipitation (IP)

IP experiments were performed as described with modifications (Chiruvella et al., 2012; Sharma et al., 2015; Totaro et al., 2011; Yeretssian et al., 2011). Protein G agarose beads (Sigma) were activated by immersing in water and incubated in IP buffer (300 mM NaCl, 20 mM Tris-HCl (pH 8.0), 0.1% NP40, 2 mM EDTA, 2 mM EGTA, and 10% glycerol) for 30 min on ice following which the beads were conjugated with appropriate antibody at 4°C for overnight to generate antibody-bead conjugate. The antibody-bead conjugates were then separated by centrifugation and incubated with rat tissue mitochondrial extracts at 4°C overnight. Then the conjugate bound to target proteins was separated and washed. The protein depletion was confirmed in the resulting supernatant by immunoblot analysis and quantified using Multi Gauge (V3.0). This immunodepleted extract was used for cleavage assay.

#### Knockdown of Endonuclease G within cells

HeLa cells (10X10^5^) were seeded in a culture petridish. 10 μg of Endonuclease G shRNA plasmid was transfected using linear/branched PEI (Polyethylenimine) polymer (Sigma, 1 mg/ml) (Longo et al., 2013; Raymond et al., 2011; Srivastava et al., 2012). Post transfection (48 h), cells were harvested, and mitochondrial extracts were prepared by differential centrifugation as described above. The knockdown was confirmed using western blotting. Cleavage assay on pDI1 following primer extension was performed using the knockdown extracts. Scrambled plasmid was also used for transfection, which acted as a control for the experiment.

#### Mitochondrial ChIP (m-ChIP)

4X10^7^ Nalm6 cells were crosslinked with 1% formaldehyde at 37°C for 15 min and quenched with 100 µl/ml of 1.375 M glycine. Mitochondria was then isolated from the cells as described above and lysed using buffer (5 mM PIPES, 85 mM KCl, 0.5% NP40). Lysed samples were then sonicated (Diagenode Bioruptor, Belgium) with 30 sec on/ 45 sec off pulse for 30 cycles. Sonicated samples were then stored at −80°C for at least 8 h (Carey et al., 2009). For immunoprecipitation, the samples were centrifuged at high speed for 15 min and the supernatant was divided for input, secondary control and experimental. Relevant antibodies (Endonuclease G or BG4) were added to the sample and allowed to bind for 8-10 h. Protein A/G-agarose beads (Sigma, USA) were added to the samples and incubated for 2 h. Samples were washed and eluted using high salt wash buffer (50 mM HEPES (pH 7.9), 500 mM NaCl, 1 mM EDTA, 0.1% SDS, 1% Triton X-100, 0.1% deoxycholate) in repeated cycles of centrifugation and then reverse cross-linked by incubating overnight at 65^°^C. Finally, DNA was purified following phenol:chloroform extraction and precipitation. The purified DNA was used for real time PCR amplification of different mitochondrial regions that either support formation of G-quadruplex structure or do not support any secondary structure formation.

#### MMEJ assay

MMEJ reactions were performed as described previously (Chiruvella et al., 2012; Sharma et al., 2015; Tadi et al., 2016). Reactions were carried out in a volume of 10 µl by incubating DNA substrates containing 9 nt microhomology (RBK7/8 and RBK27/28) with 0.5-2 µg of mitochondrial extracts in a buffer containing 50 mM Tris-HCl (pH 7.6), 20 mM MgCl2, 1 mM DTT, 1 mM ATP and 10% PEG at 30°C for 1 h. The MMEJ reactions were terminated by heat denaturation at 65°C for 20 min. The end joined products were PCR amplified using radiolabelled primer RBK13 and unlabeled primer RBK14 and the products were resolved on 8% denaturing PAGE.

#### Cloning and sequencing of end-joined junctions

MMEJ reaction products were PCR amplified and resolved on 8% denaturing PAGE. Bands of interest were cut from dried gel, resuspended in TE and 5 M NaCl and DNA was purified using phenol:chloroform extraction followed by precipitation. The purified DNA was ligated into TA vector at 16°C for 16 h and transformed into *E. coli* (DH5α). Plasmid DNA was isolated from clones of interest, digested using restriction digestion to verify the presence of insert and the positive clones were sequenced (Medauxin, India).

#### Reconstitution of 9-bp deletion

Either 2 µg of plasmid pDI1 or mitochondrial DNA was incubated with 1 µg of Endonuclease G at 37°C for 1 h. DNA was purified using phenol:chloroform extraction, precipitated, and loaded into 0.8% agarose gel. The linearized DNA was cut out of the gel and eluted. Joining reactions were performed by incubating 200 ng of eluted DNA with testicular mitochondrial extract at 30°C for 1 h. The reaction was terminated and deproteinized and used for amplifying the mitochondrial G-quadruplex forming Region I. Sample without the extract served as a control. The amplicon was ligated to TA vector and sequenced as detailed above (Medauxin, India).

#### Immunofluorescence

Immunofluorescence studies were performed as described before (Dahal et al., 2018; Kumari et al., 2019; Ray et al., 2020). Approximately 50,000 cells (HeLa, HEK293T, Rho(0), MEF or HCT) were grown in the media for 24 has described before. In the stress study, cells were either grown in the medium without FBS (serum starvation) or were irradiated (2 Gy) or treated with menadione (25 μM) for 2 h. HeLa cells were grown in chamber slides in MEM medium supplemented with 10% FBS and 1% penicillin-streptomycin (Sigma) for 24 h. The cells were stained with either 100 ng/ml of MitoTracker Red 580 (Invitrogen) or 100 ng/ml MitoTracker Green FM (Invitrogen) at 37°C in a CO2 incubator for 30 min (Tadi et al., 2016). Cells were washed twice with 1 X PBS, fixed in 2% paraformaldehyde (20 min) and permeabilized with 0.1% Triton X-100 (5 min) at room temperature. BSA (0.1%) was used for blocking (30 min) and subsequently the cells were incubated with appropriate primary antibody at room temperature (4 h). Appropriate FITC or Alexa Fluor conjugated secondary antibodies were used for detection of the signal. After washing, the cells were stained with DAPI, mounted with DABCO (Sigma) and imaged under confocal laser scanning microscope (Zeiss LSM 880 with the 63X magnifications or Olympus FLUOVIEW FV3000 with 100X magnifications). The images were processed using either Zen Lite or FV31S-SW software.

#### Gene expression analysis from TCGA data

Transcriptome data for Liver hepatocellular carcinoma (LIHC) was obtained from TCGA (https://portal.gdc.cancer.gov/) (Center, 2016). Normalized RNA-Seq by Expectation Maximization (RSEM) file was downloaded from the portal for 371 patients and 49 normal samples. Based on the TCGA barcodes, patient and normal samples were separated. Expression levels for genes like Endo G and MMEJ associated genes (Ligase III, XRCC1, MRE11A, CtIP, Exo G, Pol Q and PARP1) were extracted.

Based on the level of expression of Endo G, 371 patient samples were classified as low (n=100), medium (n=176) and high (n=95). Normal samples were also classified into low (n= 29) and high (n=20) (Fig. S15). RSEM values for MMEJ associated genes from normal and patient samples were compared in Endo G medium and high HCC patients and plotted as a Box and Whisker plot (Figure S7).

#### Statistical analysis

Every experiment was repeated multiple times with at least three biological repeats and presented. Based on the data obtained from repeats, Student’s t-test (two tailed) were performed and the error bars were calculated for the bar diagrams that represent the fold change value using GraphPad Prism ver 5.0. Error bars are shown depicting mean ± SEM (ns: not significant, *p<0.05, **p<0.005, ***p< 0.0001). Colocalization analysis in immunofluorescence assay was done using JaCoP in ImageJ software. The values were plotted in GraphPad Prism 5.0 and the significance was calculated using the same. The values were obtained for at least 50-100 cells per sample.

## Acknowledgements

We thank Mridula Nambiar, Namrata M. Nilavar, Vidya Gopalakrishnan and other members of SCR laboratory for critical reading and comments on the manuscript. We thank Monica Pandey, Rupa Kumari, Mahesh Hegde and Supriya Vartak for technical help. We also thank J. Neuzil (Australia) A Agarwal (India) and N Bhatraju for help with Rho(0) cells, S. Balasubramanian (UK) for pSANG10-3F-BG4 plasmid, Michael A. Resnick (USA) for FEN1 plasmid, Richard Baer (USA) for CtIP-pET14b plasmid, Patrick Swanson (USA) for MBP-RAG constructs, Kumar Somasundaram (India) for pIRES2-EGFP and Arun Kumar (India) for pcDNA3.1RFP. We thank Confocal, FACS and Central Animal Facilities, shRNA core facility of IISc, for their help. This work was supported by grants from CSIR (37(1692)/17/EMR-11), DAE (21/01/2016-BRNS/35074), DBT-COE (BT/PR/3458/COE/34/33/2015), IISc-DBT partnership program [BT/PR27952-INF/22/212/2018] to SCR. SD is supported by Senior Research fellowship (SRF) from IISc and HS is supported by Junior Research fellowship (JRF) from CSIR.

## Supplementary FIGURE LEGENDS

**Figure S1.**
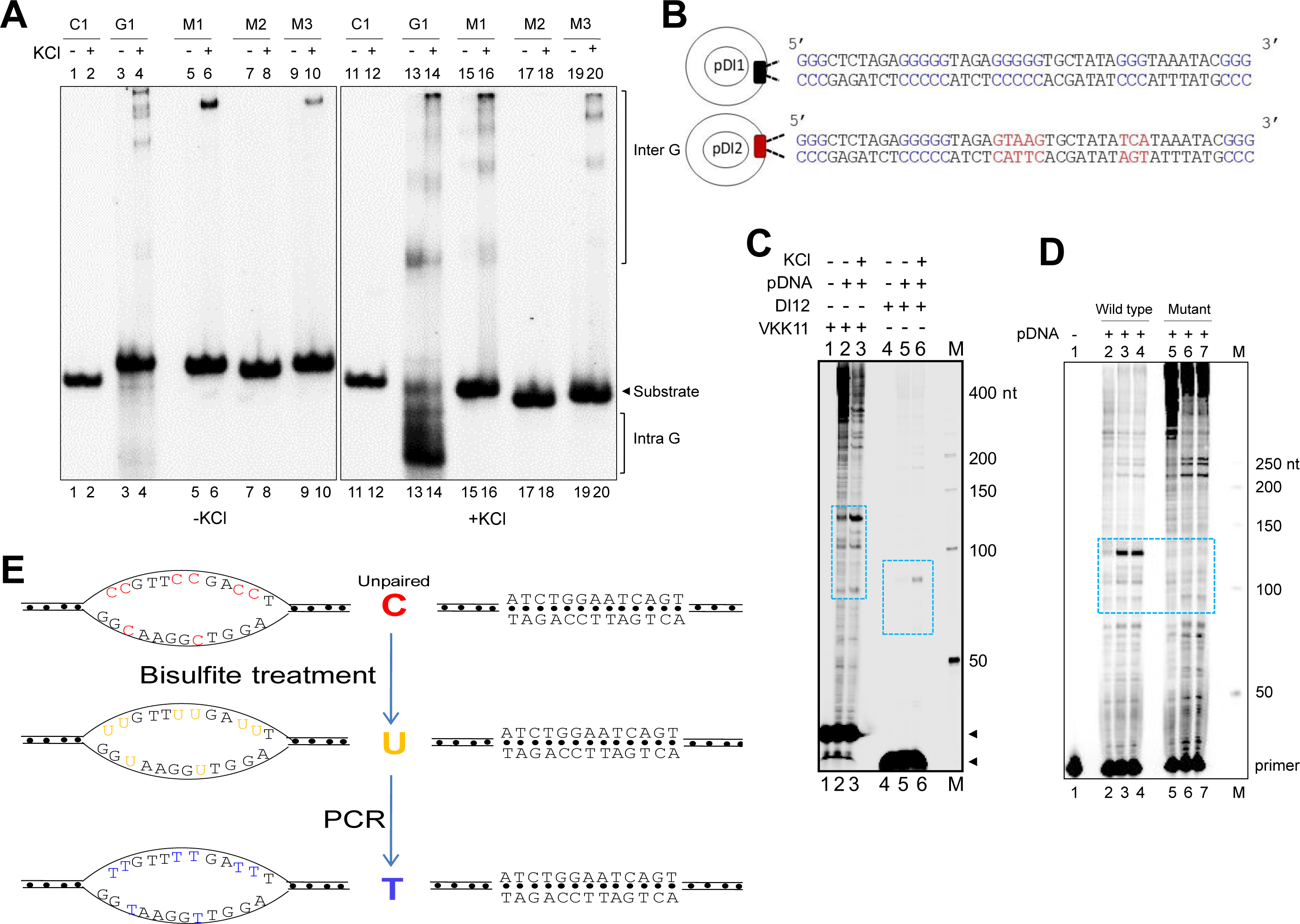
Evaluation of G-quadruplex formation at mitochondrial Region I. Related to Figure 1. **A.** Gel shift assay showing the impact of mutation of G stretches on intramolecular G4 DNA formation. Oligomeric DNA sequence is provided in Figure 1, panel A. **B.** Schematic showing cloning of the mitochondrial Region I and its mutant to generate plasmids, pDI1 and pDI2. The duplex region containing the G stretches are depicted in blue, while the mutated nucleotides are marked in red. **C.** The pDI1 plasmid, containing the mitochondrial Region I, was used for primer extension studies using radiolabeled primers (either VKK11 or pDI12) that bind at different positions of template containing the G-quadruplex-forming motif. The products were resolved on 8% denaturing PAGE. Pause sites are indicated with dotted rectangles. “M” denotes the 50 nt ladder. **D.** Plasmids, pDI1 and its mutant pDI2, containing mutation in G4 motif, were used for primer extension studies using radiolabelled primer ‘VKK11’. Pause sites are shown with dotted rectangles. “M” denotes the 50 nt ladder. **E.** Schematic showing conversion of cytosine to thymine following bisulfite modification assay. Treatment with sodium bisulfite can result in deamination of cytosine, leading to uracil when present on a single-stranded DNA. This C to U conversion can lead to C to T change following PCR and sequencing.

**Figure S2.**
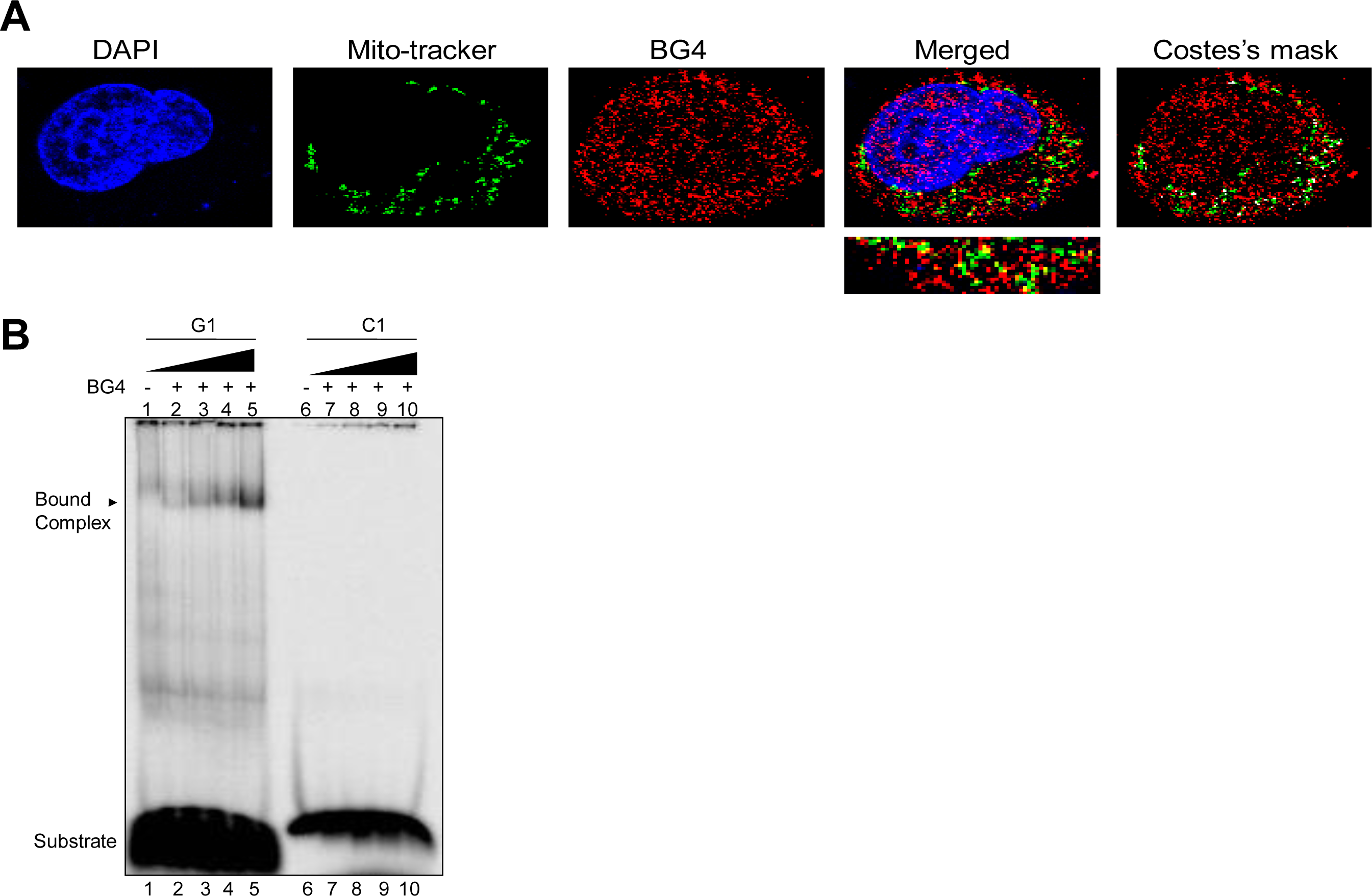
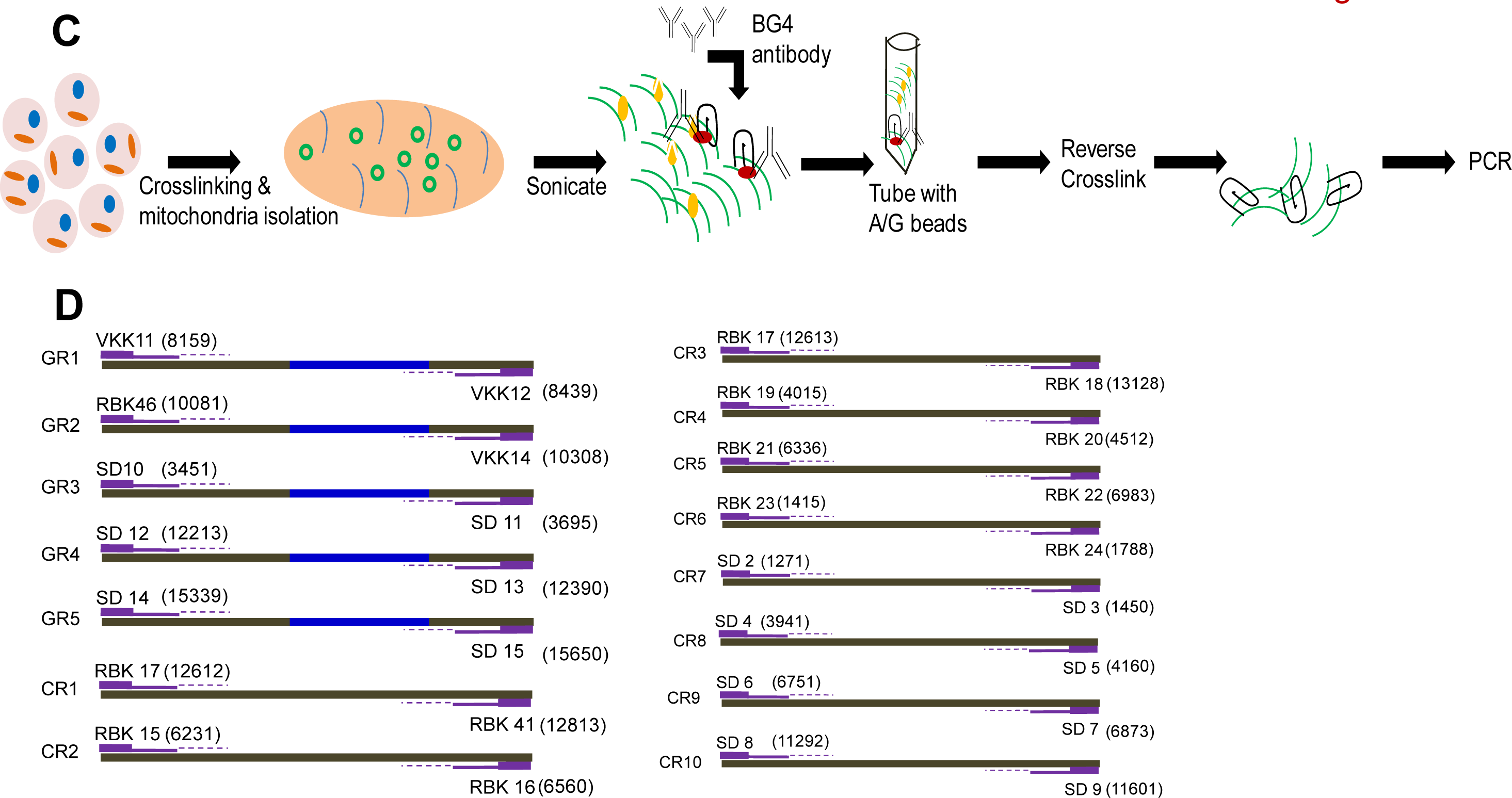
Evaluation of existence of G-quadruplex in mitochondrial DNA. Related to Figure 2. **A.** Immunofluorescence study showing colocalization of BG4 (antibody that binds to G4 DNA) and mitochondrial genome. Nucleus from HeLa cells is stained with DAPI (blue color). Mitochondria are stained with MitoTracker green FM and G4 DNA with BG4 (Alexa-Fluor 568), a merged image is shown. A merge of red and green is depicted by Coste’s mask (colocalization is represented as a white dot). The experiments were done independently in HeLa and 293T cells and the quantitation of colocalization of BG4 with MitoTracker is shown in figure 2. **B.** Increasing concentrations (0.2, 0.5, 1 and 2 µg) of purified BG4 was incubated with G-rich (G1) and C-rich (C1) oligomers and products were resolved on 5% native polyacrylamide gel. In each case a lane without protein (lane 1 and 6) served as control. The substrate and the bound complex are marked. **C.** Schematic showing the experimental strategy used for ChIP using anti-BG4. Briefly, cells were crosslinked and then mitochondria were isolated and sonicated to obtain the small fragments of mitochondrial DNA. Purified BG4 antibody was used along with protein A/G beads to pull down the BG4 bound regions. **D.** Schematic showing the position of primers used for ChIP studies. GR1-GR5 represents primers that can amplify G-quadruplex forming motifs and CR1-CR10 represents the random control regions.

**Figure S3.**
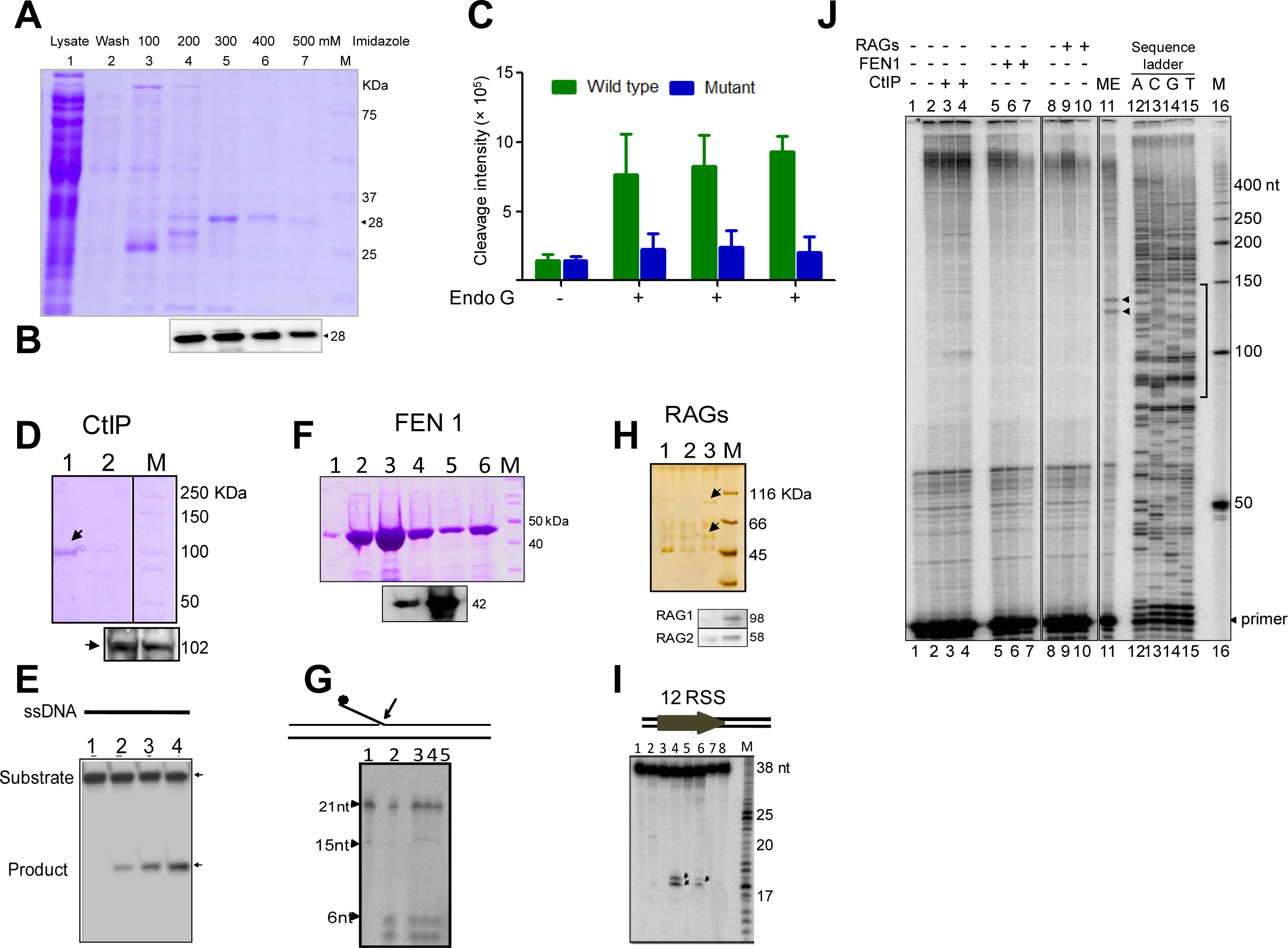
Overexpression, purification and activity assay of different endonucleases. Related to Figure 3. **A, B.**Overexpression and purification of Endonuclease G protein. The purity and identity of the protein was confirmed using SDS-PAGE (A) followed by western blotting (B). **C.** Quantification showing the efficiency of Endonuclease G mediated cleavage when a plasmid containing mutant and wild type G4 motif derived from Region I was compared. **D**. SDS gel profile showing the overexpression and purification of CtIP. Western blotting (bottom panel) was performed for confirmation of the protein. **E.** Activity assay showing resection of the substrate (ssDNA) upon the addition of increasing concentrations (30, 60 and 90 ng) of purified CtIP (lanes 2-4). **F.** SDS gel profile showing the overexpression and purification of FEN1. Western blotting confirming the identity of the protein is also shown. **G.** Activity assay showing the cleavage of the substrate with flap DNA upon the addition of increasing concentrations (50, 100, and 500 ng) of purified FEN1 (lanes 2-5). **H.** Silver stained gel profile showing the overexpression and purification of RAGs from mammalian cells. Western blotting confirming the presence of RAG1 and RAG2. **I.** Activity assay showing the cleavage of the RSS substrate upon the addition of different fractions of purified RAGs (lanes 2-8). **J.** Cleavage assay for different nucleases was performed on a plasmid containing mitochondrial Region I, pDI1. Following incubation with different purified nucleases CtIP, FEN1, and RAGs, primer extension assay was carried out using γ-^32^P radiolabeled VKK11 primer and resolved on 8% denaturing polyacrylamide gel along with sequencing ladder. Lane 1 is primer alone, lanes 2, 5, 8 are without protein, lane 11 is with mitochondrial extract and lanes 12-15 (A, C, G, T) represents the sequencing ladder. Lanes 3,4 represents (CtIP); lanes 6,7 (FEN1); lanes 9, 10 (RAGs) incubated samples. In each case, 30 and 60 ng of protein was used. ‘M’ is 50 bp ladder.

**Figure S4.**
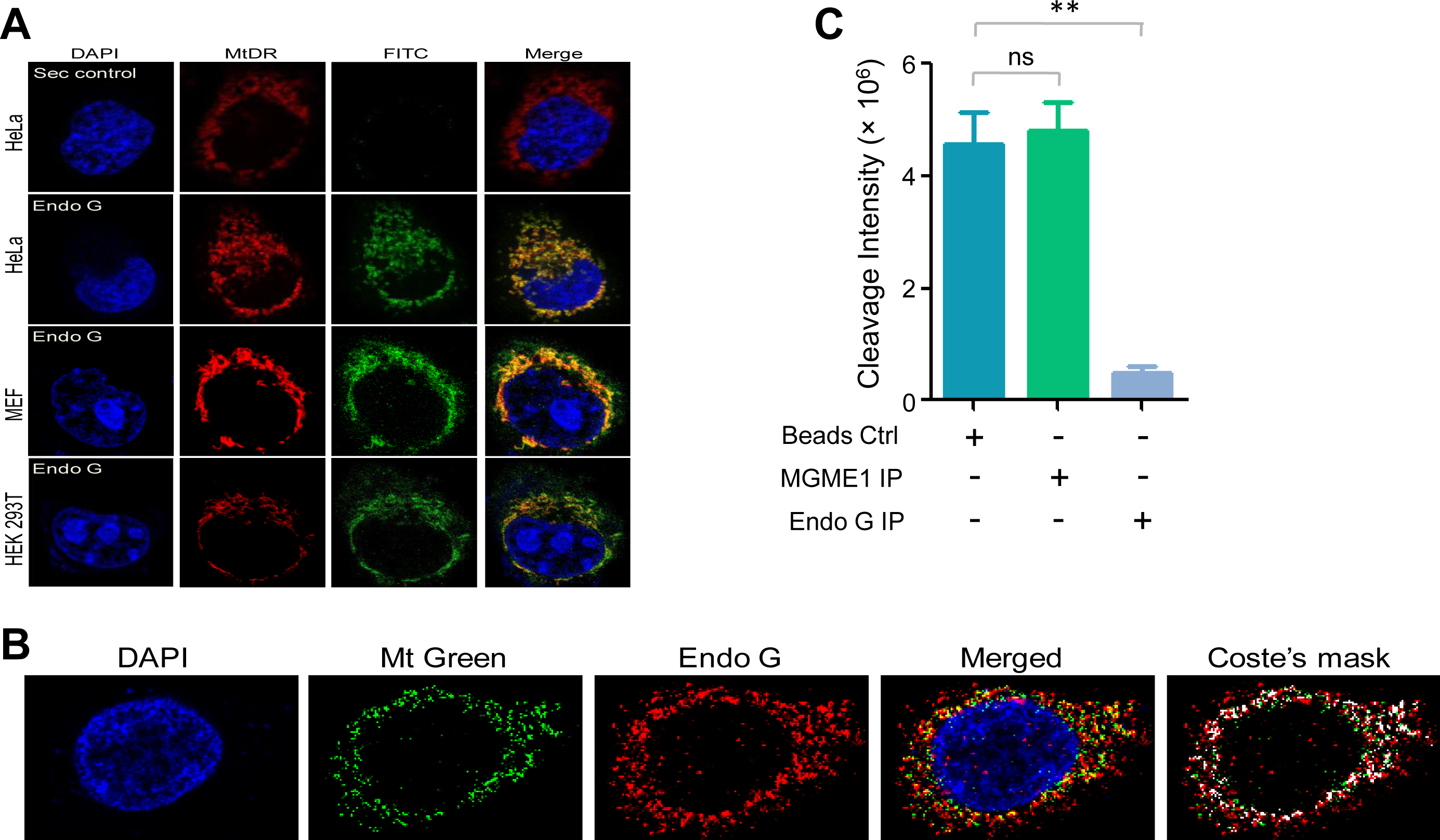
Analysis of Endonuclease G localization and function in mitochondria. Related to Figure 3. **A.** Localization of Endonuclease G in mitochondria in different cell lines. Representative images of localization of Endonuclease G to mitochondria in HeLa, MEF and HEK293T cells. FITC-conjugated secondary antibodies were used for detecting Endonuclease G proteins. MtDR is Mitotracker Red, a mitochondrial marker. DAPI is used as nuclear stain. **B.** Representative image showing colocalization of Endonuclease G using MitoTracker Green FM (Mt Green) in HeLa cells. Alexa-568 conjugated secondary antibody was used for the detection of Endonuclease G. DAPI is used to stain the nucleus. **C.** Bar diagram depicting quantitation showing the impact of immunoprecipitation of Endonuclease G based on multiple experiments is also shown.

**Figure S5.**
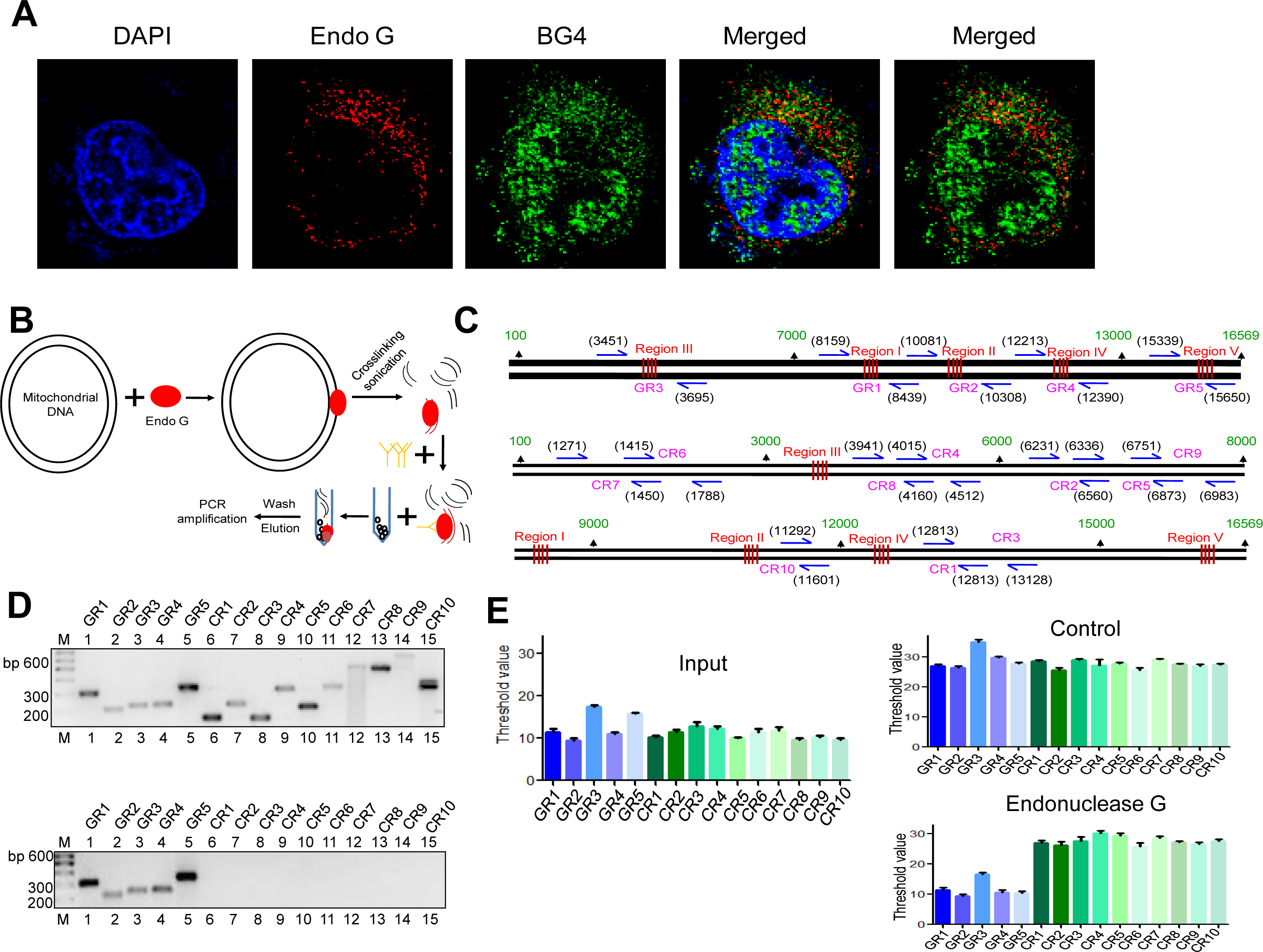
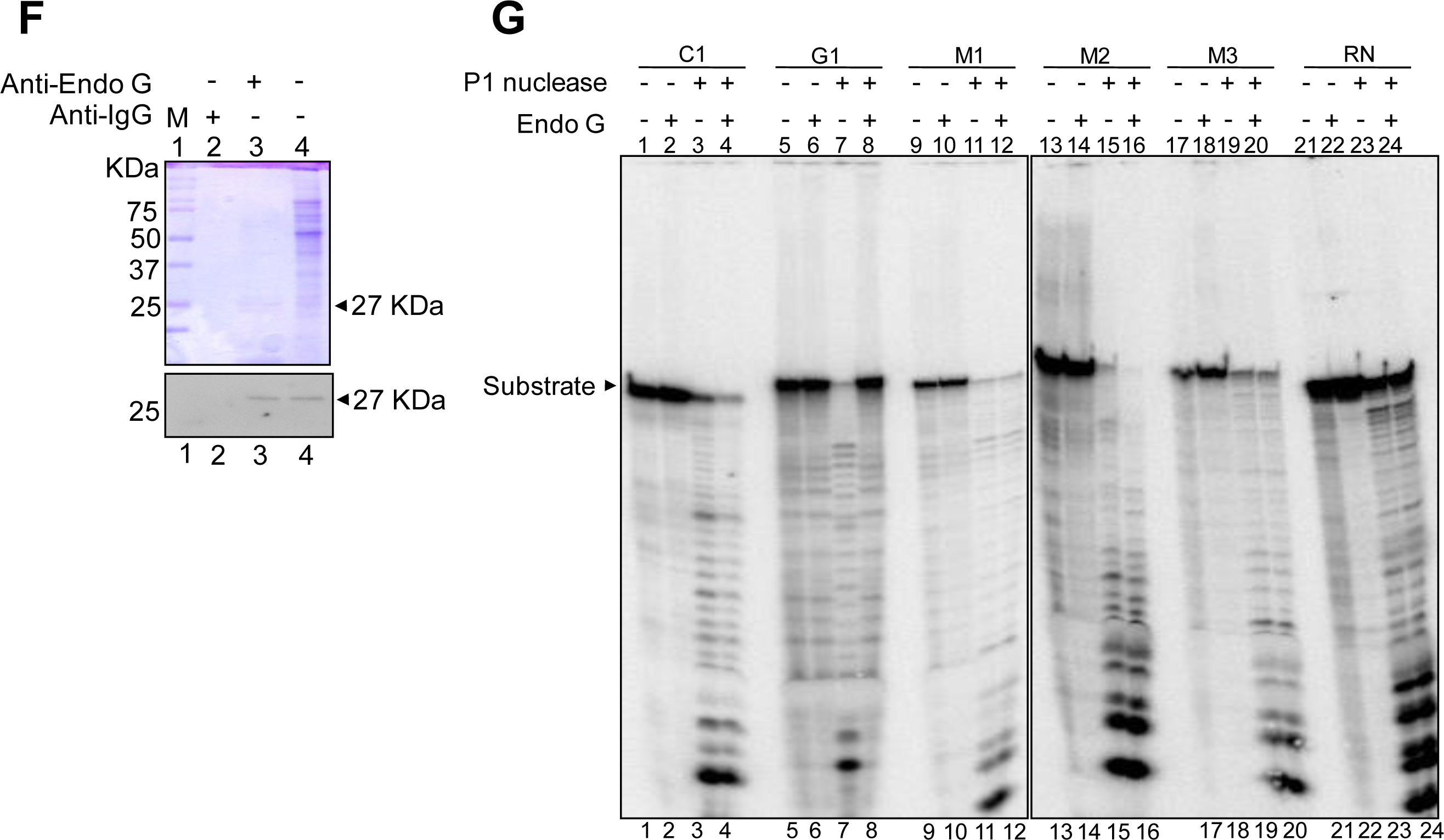
Binding of Endonuclease G to G-quadruplex regions of the mitochondrial genome. Related to Figure 4. **A.** Representative immunofluorescence images showing the colocalization of Endonuclease G and BG4. The ‘Merged’ image shown in left is a colocalization of DAPI, Endonuclease G and BG4 foci, while ‘Merged’ in right shows colocalization of Endonuclease and BG4 foci. **B.** Schematic showing the binding of purified Endonuclease G to the mitochondrial genome and the pull-down of a bound region using Endonuclease G antibody and protein A/G beads. These regions were then used for semi-quantitative and real-time PCR using appropriate primers. **C.** Schematic showing the position of primers used for ChIP studies. The upper panel shows the primer positions for G-quadruplex forming regions and lower panels for control regions. **D.** 5 G-quadruplex forming regions and 10 random regions were used for amplification using m-ChIP DNA as template (lower panel). Upper panel shows the amplification for input DNA. **E.** Bar graph was plotted for the threshold cycle against different primers following real-time PCR. Input DNA served as template control. ‘Antibody’ control in which no antibody was added served as negative control. Blue bars represent G-quadruplex regions, while green bars represent the control regions. Error bar represents three independent biological repeats. **F.** SDS profile and western blotting of the pulldown sample when mitochondrial DNA was incubated with mitochondrial extracts. For other details refer main Figure 4. **G.** P1 nuclease footprinting to demonstrate the binding of Endonuclease G to G-quadruplex forming Region I. Radiolabelled oligomers were incubated with Endonuclease G protein and subjected to P1 nuclease and electrophoresed on 18% denaturing PAGE. In each case, lanes 1, 5, 9, 13, 17, 21 are substrate alone, lanes 2, 6, 10, 14, 18, 22 are Endonuclease G alone, lanes 3, 7, 11, 15, 19, 23 are P1 nuclease alone treated samples and lanes 4, 8, 12, 16, 20, 24 are Endonuclease G plus P1 nuclease treated samples. C1 is C-strand, G1 is G strand, M1, M2 and M3 are mutants while random sequence (RN) is the oligomer with equal G-C content as G1 in a random manner. 50 ng of purified Endonuclease G and 0.03 U of P1 nuclease was used for the assay.

**Figure S6.**
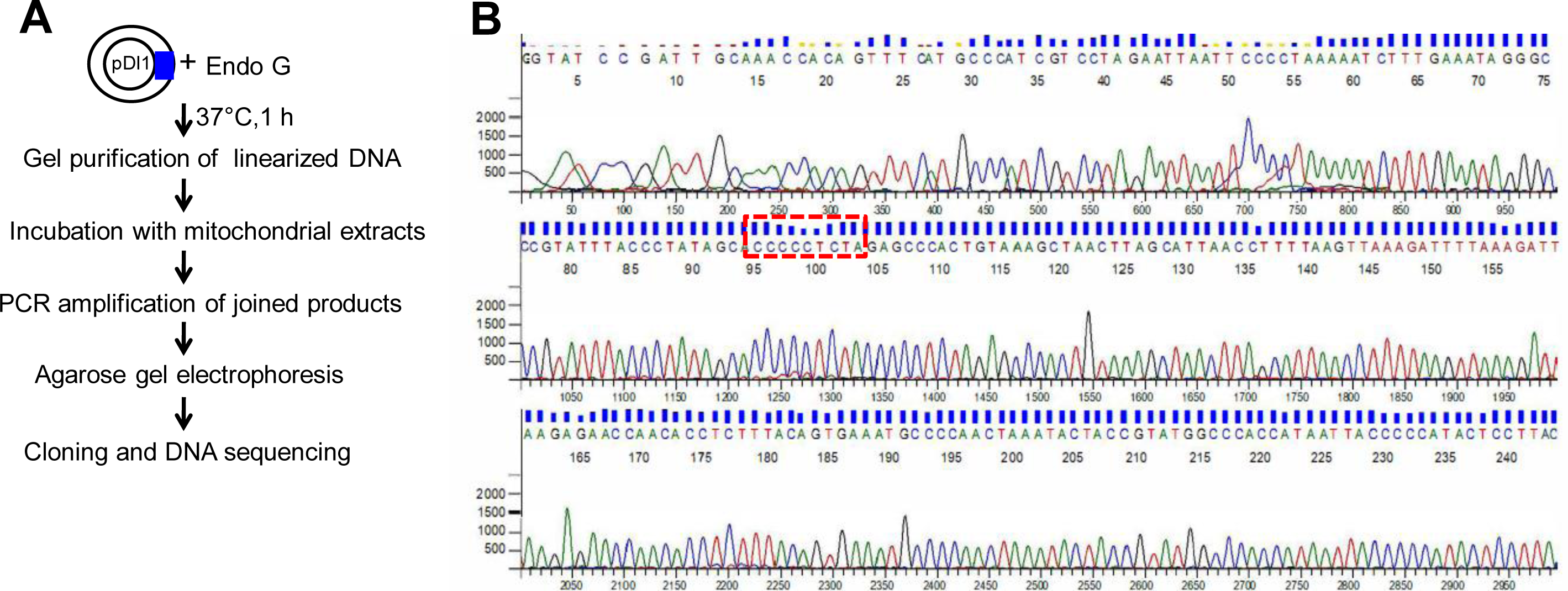
Chromatogram showing sequence of the representative clone with ‘9 bp deletion’ following the reconstitution of the whole process. Related to Figure 5. **A.** Strategy used for reconstitution of mtDNA deletion, when the Region I was present on a plasmid DNA. See also Figure 6. **B.** Representative sequence of a clone showing recapitulation of ‘9 bp deletion’. In this case a 9 bp direct repeat was deleted when the mitochondrial genome was treated with purified Endonuclease G followed by DSB repair using mitochondrial extracts.

**Figure S7.**
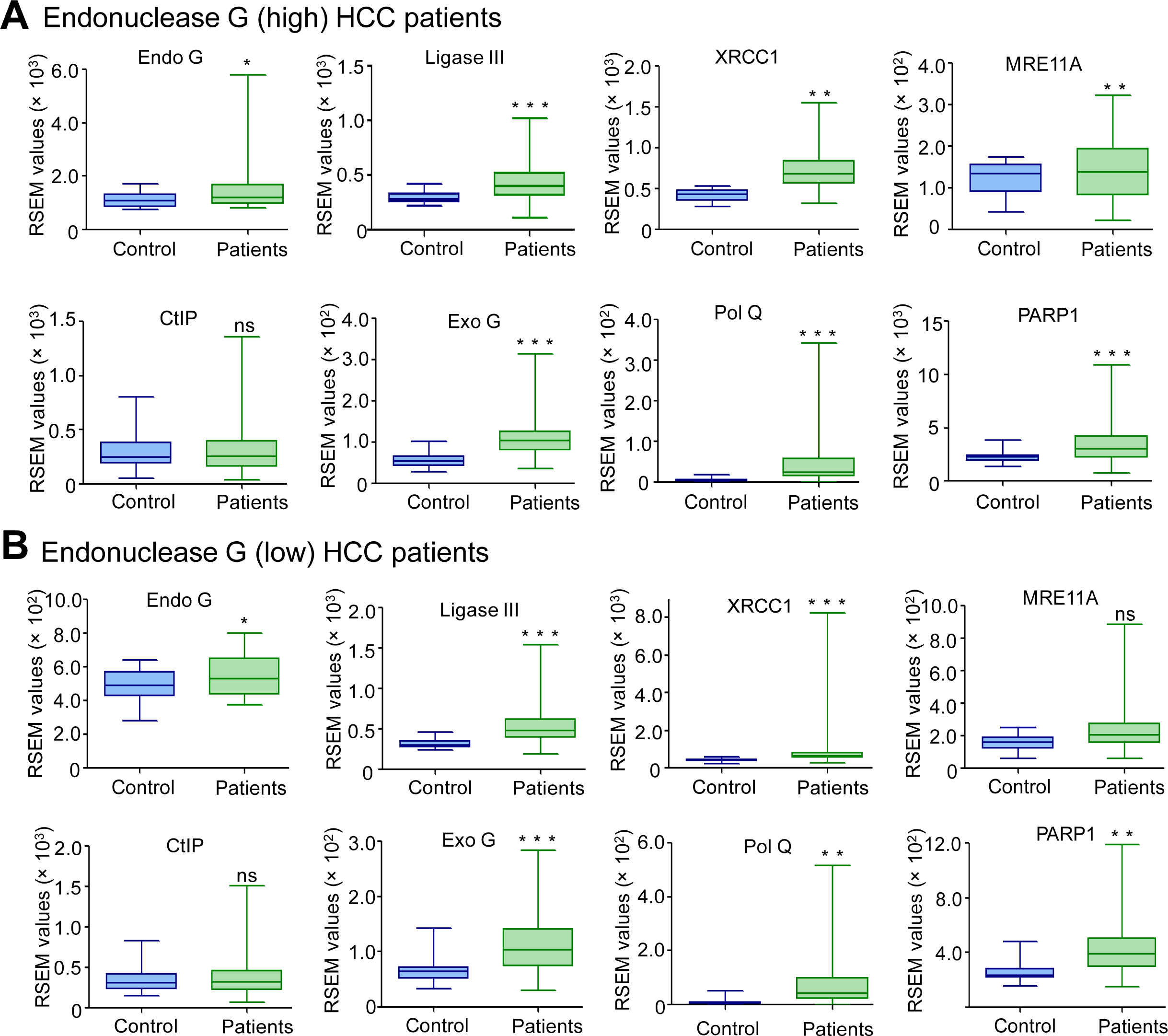
Expression of Endo G and MMEJ associated genes in hepatocellular carcinoma patients. Related to Figure 6. **A.** Box and Whisker plot showing the expression of Endo G and MMEJ related genes (Ligase III, XRCC1, MRE11A, CtIP, Exo G, Pol Q, PARP1) in the Endo G high patient samples (n=95) compared to that of normal (n=20). **B.** Box and Whisker plot showing the expression of Endo G and MMEJ genes in HCC patient samples with medium expression of Endo G compared to normal samples. In all panels, Y-axis depicts the RSEM values, while X-axis depicts the type of sample.

